# Evolutionary origin and demographic trajectory of tomato spotted wilt virus (*Orthotospovirus tomatomaculae*)

**DOI:** 10.1101/2025.03.10.642482

**Authors:** José A. Castillo, Lenin Ramirez-Cando

## Abstract

Tomato spotted wilt orthotospovirus (TSWV) is a harmful pathogen that causes severe disease in tomato, pepper, and other horticultural and agronomic crops. The genome comprises three linear ssRNA molecules (segments L, M, and S) unequally packed in nucleocapsids. Although the genome structure and molecular mechanism of infection are well understood, the evolutionary dynamics require further analysis to know what we seek in this study: to determine the most probable viral ancestor, the date of divergence, the ancestral geographical source of dispersal, and the demographic history of TSWV. We employed the Bayesian framework to analyze whole genome sequences of 136 isolates of TSWV obtained from different sites worldwide and with a sampling window of 35 years. In independent analyses, we performed phylodynamic estimations with segment L, M, and S sequence data. Our results showed that the clock rates of different genome segments ranged from 1.678×10-4 to 2.444×10-4 substitutions/site/year. Likewise, the time to the most recent common ancestor shows variations according to the dataset used for the inferences; however, the most probable date of TSWV divergence is around 1768 CE. Our phylogeographic exploration showed concordant results for the three genomic segments analyzed; that is, the TSWV population originated in South Korea and, from there, early expanded to Europe and then to North America and other continents. Past population dynamics analysis showed that the virus experienced two major population expansions that coincided with the increase of the agricultural frontier and the emergence of new species of insect vectors.

## Introduction

Tomato spotted wilt virus (TSWV) is a viral plant pathogen that causes severe losses in crop production. It can infect up to 1000 plant species from more than 90 plant families, including economically important crops such as tomato, pepper, peanut, and others (Parrella et al., 2003; Oliver & Whitfield, 2016). TSWV is considered a severe pathogen for crops. In tomatoes, the yield can decrease from 38.7% to 100% in severe outbreaks, especially in early infection stages (Fiederow & Kralowska, 1995; Pappu et al., 2009). In other crops, the losses may range from 50 to 90% in lettuce, 5 to 80% in peanuts, and more than 70% in peppers (EPPO, 2024). Usually, plants infected with TSWV show foliage covered with spots, and the leaves turn brown, roll, and wilt. The fruits with disease symptoms appear bronzed with yellow or necrotic ringspots and streaks with mosaic and concentric ring patterns (Kenyon et al., 2014).

TSWV, also known as *Orthotospovirus tomatomaculae*, is classified in the order Bunyavirales, family Tospoviridae that typically harbors segmented single-strand RNA (ssRNA) genome. The genome comprises three different ssRNA fragments that encode a few protein-coding genes and are replicated following the mechanism of negative strand viruses. However, some TSWV RNAs can be ambisense; that is, both direct and complementary sequences are used to make gene products. The middle (M) and small (S) segments encode two genes each in opposite or ambisense polarity: the segment M, the NSm (Movement protein) and Glycoprotein precursor proteins, and the segment S, the NSs (Silencing suppressor) and the N (Nucleocapsid) proteins. The large (L) segment encodes the multipotent enzyme called RNA-dependent RNA polymerase (RdRps) that transcribes and replicates the viral genome in the host cell (Oliver & Whitfield, 2016). A feature common to TSWV is the widespread exchange of genetic material between viruses of the same species or other related species. The genetic exchange can occur naturally through recombination or reassortment when two or more virus lineages co-infect the same host plant (Butković et al., 2021).

TSWV is spread through insect vectors. Thrips (mostly *T. tabaci*) were the traditional vectors of TSWV (Jones, 2005), but later, different species of *Frankliniella* (usually known as western flower thrips) arose as the most common and prevalent vectors of TSWV (Wan et al., 2020), *Frankliniella occidentalis* being the most effective vector (Prins & Goldbach 1998). The geographic distribution of the TSWV and its thrips vector is often coincident (Jones, 2005).

TSWV has dispersed throughout different countries since the 1930s; however, the first report of its existence dates back to 1915 (Brittlebank, 1919). The dynamics of the infection over the years have been irregular, with alternating periods of severe disease or reduction of symptoms. Control of the disease is often associated with programs to eradicate insect vectors. The use of resistant varieties has also been successful in avoiding this pathogen. Tomato and pepper cultivars carrying the resistance gene Sw-5b or Tsw, respectively, are well-known and widely used (De Oliveira et al., 2018; Moury et al., 1997). These resistant cultivars have been one of the most effective TSWV management strategies for years; however, lately, the emergence of novel resistance-breaking strains of TSWV worldwide has been seen (Chinnaiah et al., 2023).

The tomato spotted wilt disease was first identified in the year mentioned above, 1915 (Brittlebank, 1919). Later, Samuel & collaborators (1930) demonstrated that this disease is caused by a virus named as we know it today (Oliver & Whitfield, 2016). Although TSWV has been identified for almost 100 years, little is known about its evolutionary origin, which may help develop successful and sustainable management strategies for this pathogen. Using phylogeny-based methods, we sought to understand the evolutive origin and spread of the TSWV population in this work. In particular, we focused on addressing questions about the possible location where the TSWV emergence occurred and the time since it separated from its most recent ancestor. We also sought to identify the most recent ancestor and population dynamics of TSWV throughout its short evolutionary history. The recent availability of a large number of genomic sequences and the development of robust methods for phylogenetic reconstruction and testing evolutionary hypotheses allowed us to estimate with a high degree of confidence different evolutive parameters that may help to understand better the biology and the evolutionary process followed by this important pathogen.

## Results

### TSWV sequences and recombination analysis

The whole genome sequences of TSWV were obtained from databases and were aligned to assemble three different datasets that correspond to the three RNA segments of the tripartite TSWV genome. All sequence alignments share the same isolates of TSWV; therefore, in this work, we analyzed a dataset of 136 TSWV viruses (see Table S1). The sequence alignment of segment L consists of 9029 nt, and the alignment of segments M and S contains 5077 and 3290 nt, respectively. We performed recombination analyses on the three datasets, which allowed us to find numerous TSWVs that presumably experienced recombination. We discovered 17 virus isolates that were identified as recombinants, and we further identified the site where recombination presumably occurred (the breaking points, see Table S2). The major recombination sites found in the segment L aligned sequences are in the coding region of the RdRp gene (RNA-dependent RNA Polymerase), which is the only gene of this RNA. The most abundant recombination breakpoints in the segment M sequences correspond to the GnGc coding region in isolates from China, South Korea, the USA, and Italy. Similarly, the common recombination sites found in the segment S sequences were in a region that corresponds to the nonstructural protein NSs in isolates from Turkey, Italy, the USA, and Australia (Table S2). We removed these recombinants from our datasets for subsequent phylogenetic analyses.

We estimated the suitability of the TSWV sequence dataset for performing phylogenetic analyses by calculating the index of substitution saturation (*Iss*). Results indicate that TSWV sequences are proper for phylogenetic reconstruction. The calculated *Iss* of segments L (*Iss*_*L*_= 0.069-0.072), M (*Iss*_*M*_= 0.099-0.103) and S (*Iss*_*S*_= 0.046-0.049) are significantly lower than the respective critical value of the index of substitution saturation *Iss*.*c* (*Iss*.*c*_*L*_= 0.816-0.855; *Iss*.*c*_*M*_= 0.811-0.851; *Iss*.*c*_*S*_= 0.73-0.811), indicating that aligned sequences have not experienced substitution saturation and consequently contain sufficient phylogenetic information.

### Temporal signal evaluation

We sought temporal signal evidence in the three TSWV sequence datasets. Using the BETS test, we calculated the ratio of the log marginal likelihoods (known as the logBF) of the two models, the heterochronous and the isochronous. Results support the heterochronous model for segments L, M, and S sequence data (see Table 1). To confirm the clock-like behavior of segments L and M sequence data, we performed the date randomization test in which we randomly shuffled the tip dates among the genomes in the dataset ten times and compared the clock rate estimates between the randomized models and the actual dataset. After running the analyses in BEAST, we verified that the ten randomized datasets and the real dataset reached an ESS of over 200 for all parameters when using the constant size and strict clock settings. According to Duchene et al., 2015, two criteria are proposed to determine whether there is a sufficient temporal signal: i) the mean rate estimated with the actual dataset does not overlap with any of the 95% credible intervals of those estimated from the date-randomized data sets; ii) the mean and 95% credible interval of the original rate estimate does not coincide with respective values of those from the date-randomized. As we can see in Figure 1, the results show that there is no overlap between the mean and the 95% credible intervals of the original rate estimate and any of those from the date-randomized data set, thus fulfilling the strictest criteria that confirm the sequence data contain temporal signal to calculate evolutive rates and estimate the tMRCA.

**Table 1.** Results of Bayesian evaluation of temporal signal of TWSV genomic sequence data. The path sampling (PS) method was employed to calculate marginal likelihood (values represent the media of three independent analyses). The Bayes factor was calculated as the ratio of the sequences with heterochronous to isochronous time data.

| Genome segment | Temporal data models | Marginal likelihood (log) | Bayes factor (log) |
| --- | --- | --- | --- |
| Segment L | Heterochronous | -51259.602089135 | 11.14 |
|  | Isochronous | -51270.7447686369 |  |
| Segment M | Heterochronous | -30067.0122366973 | 5.98 |
|  | Isochronous | -30072.9934058137 |  |
| Segment S | Heterochronous | -18863.1723737887 | 3.06 |
|  | Isochronous | -18866.2288628738 |  |

**Figure 1.**
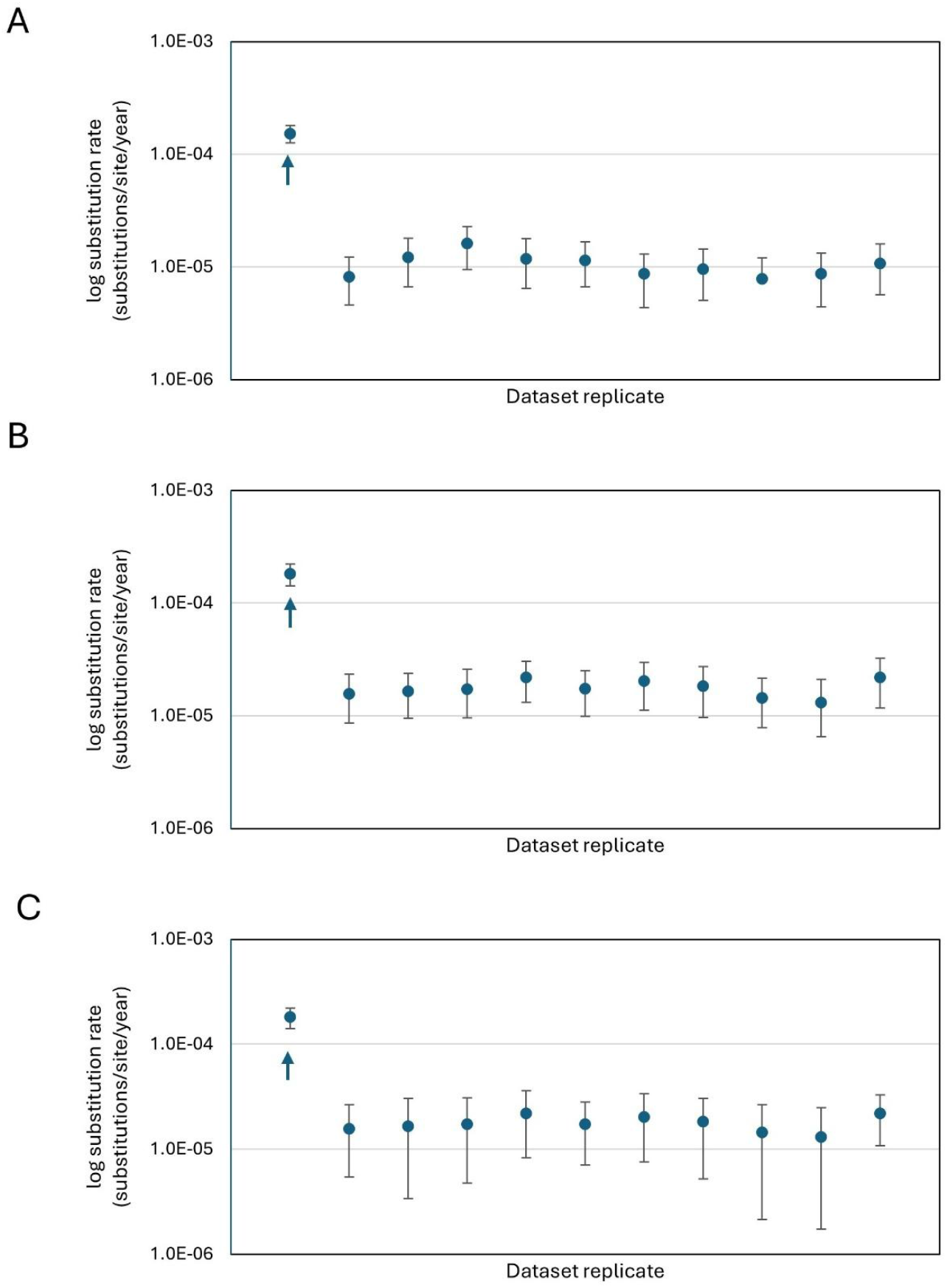
The date randomization test compares the clock rate calculated with the actual dataset of A) Segment L, B) Segment M, and C) Segment S and with a dataset where sampling times were randomly shuffled. The dots represent the mean value for the rate parameter, and the error bars represent the rate parameter’s 95% HPD interval. The arrow refers to the actual model with true tip dates.

### Evolutionary rates and timescales

Once confirmed the temporal signal is sufficient to be able to estimate the evolutionary rate, we run analyses to test different clock and tree models (i.e., strict or relaxed clock models versus coalescent constant size, exponential growth, or coalescent Bayesian Skyline demographic models). We quickly reached convergence with a few MCMC chains (30 million) when testing the combination of strict clock and constant size or exponential growth. The opposite was true when testing the relaxed clock model with the tree models since this combination caused slower convergence for all three sequence datasets. The coefficient of variation indicates whether the data set fits better to a strict or a relaxed clock. All three datasets (i.e., segments L, M, and S sequence data) reach a coefficient of variation around 0.4-0.6 depending on the tree model accompanying the analysis. According to Drummond and Bouckaert (2015), values closer to zero indicate the data are more clock-like, and a strict clock is more appropriate. There are no established values that define the upper and lower limits of the coefficient of variation to decide whether a dataset fits better to a strict or relaxed clock model. In the TSWV case, although the value of the coefficient of variation does not approach zero, it is nevertheless not close to or greater than 1, indicating that any of the clock models could describe the data. In the face of this uncertainty, we calculate the marginal likelihood of strict and relaxed clock models in combination with the three tree models stated above to define through marginal likelihood which are the best models for the data.

Table 2 shows the results of the marginal likelihood of all clock and tree model combinations tested. All three segments’ datasets are better described by the relaxed clock model accompanied by the coalescent Skyline tree model. However, it remains possible that a speciation model may have better performance for the TSWV sequences. Therefore, we tested the Birth-Death Skyline prior (BDSKY, Stadler et al., 2013) combined with the relaxed clock. The relaxed clock-BDSKY model combination had a lower marginal likelihood value than the coalescent Skyline counterpart (Table 2), which looks reasonable considering that the sequence dataset comes from a single virus species.

**Table 2.** Model comparison for TSWV datasets. Marginal likelihood) estimates from the combination of two clock and three tree models applied to segments of the TSWV genome. Marginal likelihood values were calculated using the path sampling method and are ordered from highest to lowest for each genome segment.

| Genome segment | Molecular clock | Tree prior | Marginal likelihood (log) | Bayes factor (log) |
| --- | --- | --- | --- | --- |
| Segment L | Relaxed | Coalescent Skyline | -18863.1723737887 |  |
|  | Relaxed | Birth-Death Skyline | -18865.6338936905 | 2.46 |
|  | Strict | Coalescent Skyline | -18889.8804612274 | 24.25 |
|  | Relaxed | Constant size | -18896.9032173932 | 7.02 |
|  | Relaxed | Exponential growth | -18910.2836853131 | 13.38 |
|  | Strict | Exponential growth | -18942.9576753720 | 32.67 |
|  | Strict | Constant size | -18947.2639055577 | 4.31 |
| Segment M | Relaxed | Coalescent Skyline | -30067.01223669 |  |
|  | Relaxed | Birth-Death Skyline | -30081.34763996 | 14.34 |
|  | Relaxed | Constant size | -30097.61223187 | 16.26 |
|  | Relaxed | Exponential growth | -30113.58465634 | 15.97 |
|  | Strict | Coalescent Skyline | -30114.72253402 | 1.14 |
|  | Strict | Constant size | -30149.10418739 | 34.38 |
|  | Strict | Exponential growth | -30156.65546450 | 7.55 |
| Segment S | Relaxed | Coalescent Skyline | -51259.602089135 |  |
|  | Relaxed | Birth-Death Skyline | -51274.114181637 | 14.51 |
|  | Relaxed | Coalescent Skyline | -51280.299651773 | 6.19 |
|  | Relaxed | Constant size | -51288.257194238 | 7.96 |
|  | Strict | Exponential growth | -51403.845122629 | 115.59 |
|  | Strict | Exponential growth | -51433.628398090 | 29.78 |
|  | Strict | Constant size | -51439.254745664 | 5.63 |

Having already defined the best models for the TSWV sequence data, we can then extract the relevant information. The clock rate of TSWV according to sequence analyses of segments L, M, and S are depicted in Table 3. The clock rate (ORCRates) values differ depending on the genomic segment analyzed; however, this variation is minor since it only differs by ∼1×10^−4^ (between the slowest and fastest rate of segments L and M, respectively, see Table 3). Similarly, the estimated date of TSWV emergence varies according to the genomic segment analyzed (Table 3). It is around the end of the 18th century (1768 CE, according to segment L data) and the beginning of the 19th century (1812 or 1843 CE in concordance with segments M and S sequences, respectively).

**Table 3.**
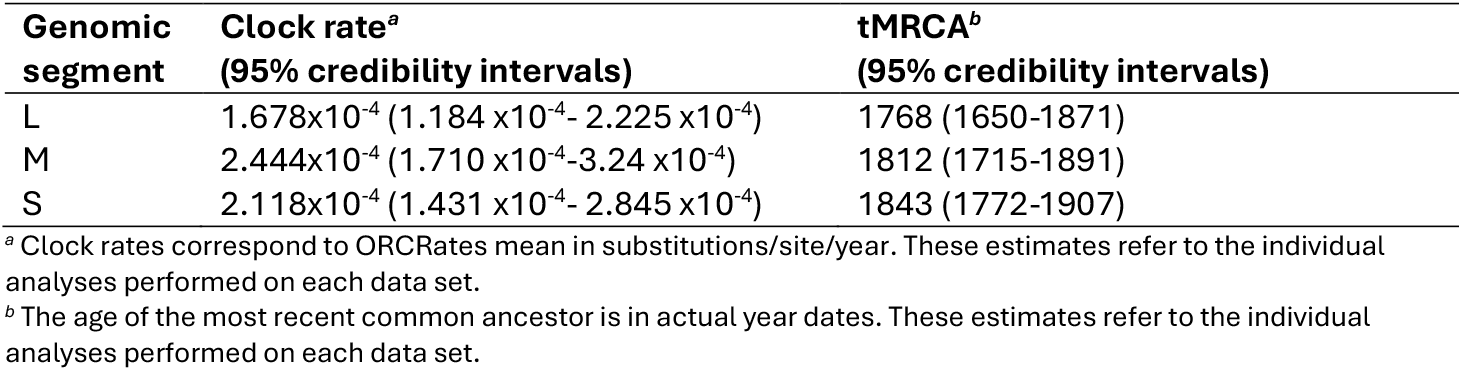
Clock rates and time to the most recent common ancestor of TSWV genomic segments.

| Genomic segment | Clock rate<sup>a</sup><br>(95% credibility intervals) | tMRCA<sup>b</sup><br>(95% credibility intervals) |
| --- | --- | --- |
| L | 1.678x10<sup>-4</sup> (1.184 x10<sup>-4</sup>- 2.225 x10<sup>-4</sup>) | 1768 (1650-1871) |
| M | 2.444x10<sup>-4</sup> (1.710 x10<sup>-4</sup>-3.24 x10<sup>-4</sup>) | 1812 (1715-1891) |
| S | 2.118x10<sup>-4</sup> (1.431 x10<sup>-4</sup>- 2.845 x10<sup>-4</sup>) | 1843 (1772-1907) |

<sup>a</sup> Clock rates correspond to ORCRates mean in substitutions/site/year. These estimates refer to the individual analyses performed on each data set.
<sup>b</sup> The age of the most recent common ancestor is in actual year dates. These estimates refer to the individual analyses performed on each data set.

### Phylogenomic trees

Bayesian maximum clade credibility (MCC) trees were inferred with the best combination of clock and tree models calculated for segments L, M, and S genomic sequences (see Table 2). The MCC trees are depicted in Figure 2. The topology of the L, M, and S trees shows substantial differences among them. Figure S1 makes it easy to visualize the similarities and differences between the trees. We calculated the Robinson–Foulds (RF) distance between these trees using branch length values. The RF measure of the pairwise comparisons of the L–M, M-S, and L-S trees show low coincidence, which is 25%, 28%, and 29%, respectively. We also used distance matrices of the L, M, and S phylogenies to compare trees by calculating Euclidean pairwise distances. The estimated distances for the L-M, M-S, and L-S pairs are considerable and exceed the value of 3000 (L-M=3309; M-S=3467 and L-S=3340). Despite the differences, it is still possible to distinguish three major clades on trees. Clade I is composed mainly of isolates from Europe that infect mostly tomato, secondary pepper, and other crops. Clade II contains isolates from Asia (China and South Korea), infecting mainly pepper and other plant species. Clade III comprises isolates from different locations, including Africa (South Africa and others), North America (USA, Mexico), Australia, and Asia.

**Figure 2.**
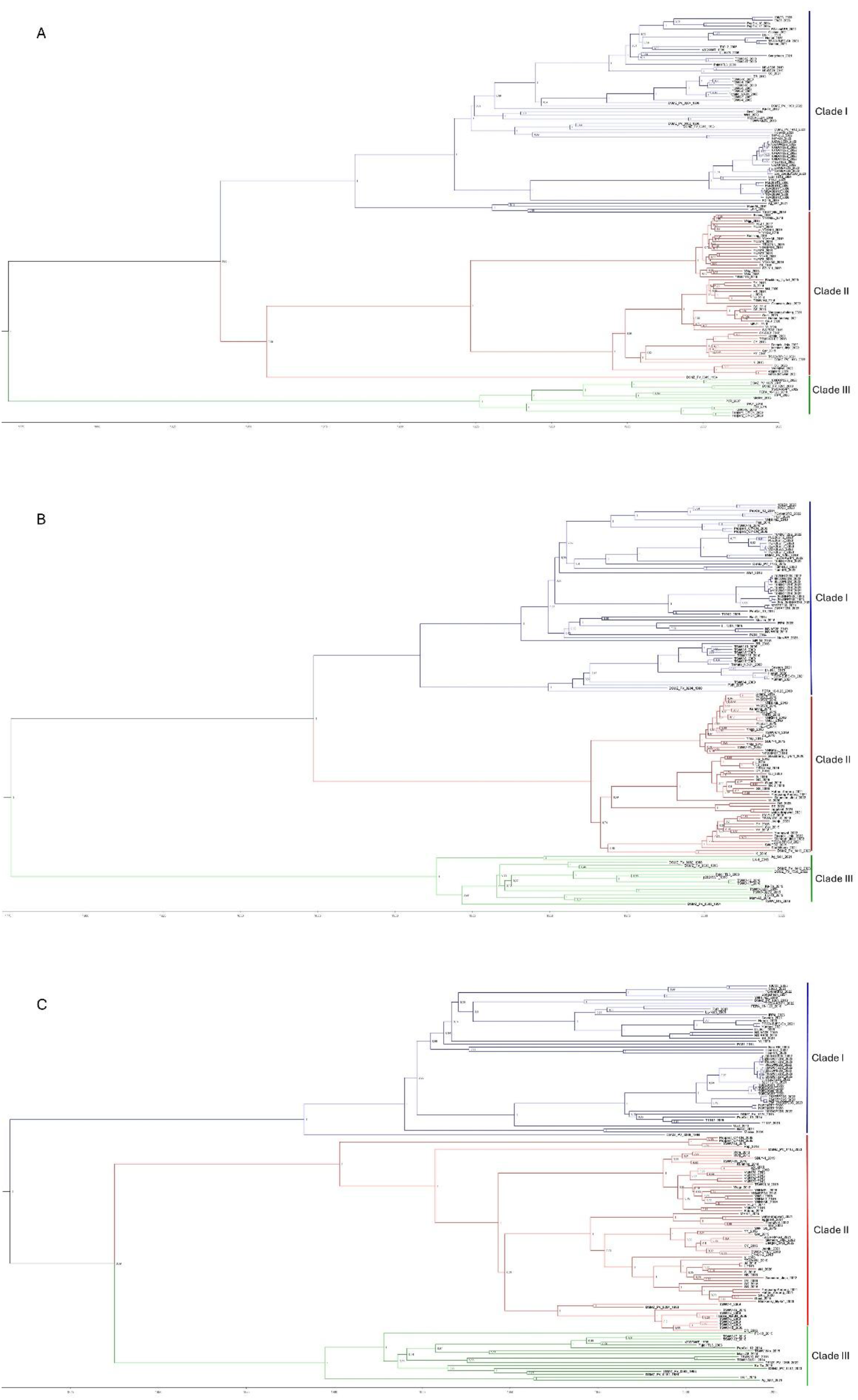
Time-scaled maximum clade credibility trees of TSWV inferred from whole-genome sequence data of A) segment L, B) segment M, and C) segment S. The color of tree branches and nodes represent major clades. Tree nodes show posterior probability values, and branch lengths are scaled according to the time indicated in the horizontal axis. Trees were estimated using the Bayesian-MCMC method with relaxed clock and Skyline models, and FigTree was used for visualization.

### Phylogeography and Demographic History

We investigated the geographical pattern of TSWV spreading around the globe, including its possible place of origin. We examined the diffusion patterns and rates among localities, considering that the virus could migrate across discrete geographical locations. We performed the phylogeography for the TSWV genome segments in separate analyses. Figure 3 shows the spatial dispersion of TSWV according to the different genomic segments’ datasets. Results related to geographical origin are coincident with phylogeographical analyses of segments L, M, and S data, indicating that the TSWV place of origin is South Korea, from where it has spread to other localities. Initially, European countries were the main recipients of TSWV from South Korea, and later, they became a secondary focus of dispersion. It is worth mentioning that the European countries where TSWV arrived are slightly different according to the genomic segment analyzed. Bulgaria and Italy were the first recipient countries according to segment M and S and Bulgaria and Spain, consistent with segment L results. According to our estimates, TSWV would have arrived in the USA between 1930 and 1935, from where it spread to Mexico and other countries. Although the origin of the virus (South Korea) is clearly established and is concordant among the findings of segments L, M, and S analyses, the pattern of dispersion slightly varies according to the genomic segments used for the phylogeographic dispersion. For example, TSWV reached African countries (South Africa and Zimbabwe) from Korea through Europe according to phylogeographic analysis performed with the segment M. However, the inference with segments L and S shows that the migration pathway was directly from Korea to South African countries. Similarly, there are differences in spreading pathways to India (Bangalore), Australia, Iran, and a few other countries (see Figure 3). We estimated our Bayesian phylogeographic analyses’ statistical support by calculating the migration pathways’ BF. Figure S2 shows that most BFs exceed the value of 2 or 6, indicating positive or strong support, respectively (Kass & Raftery, 1995) for TSWV migration between the indicated countries.

**Figure 3.**
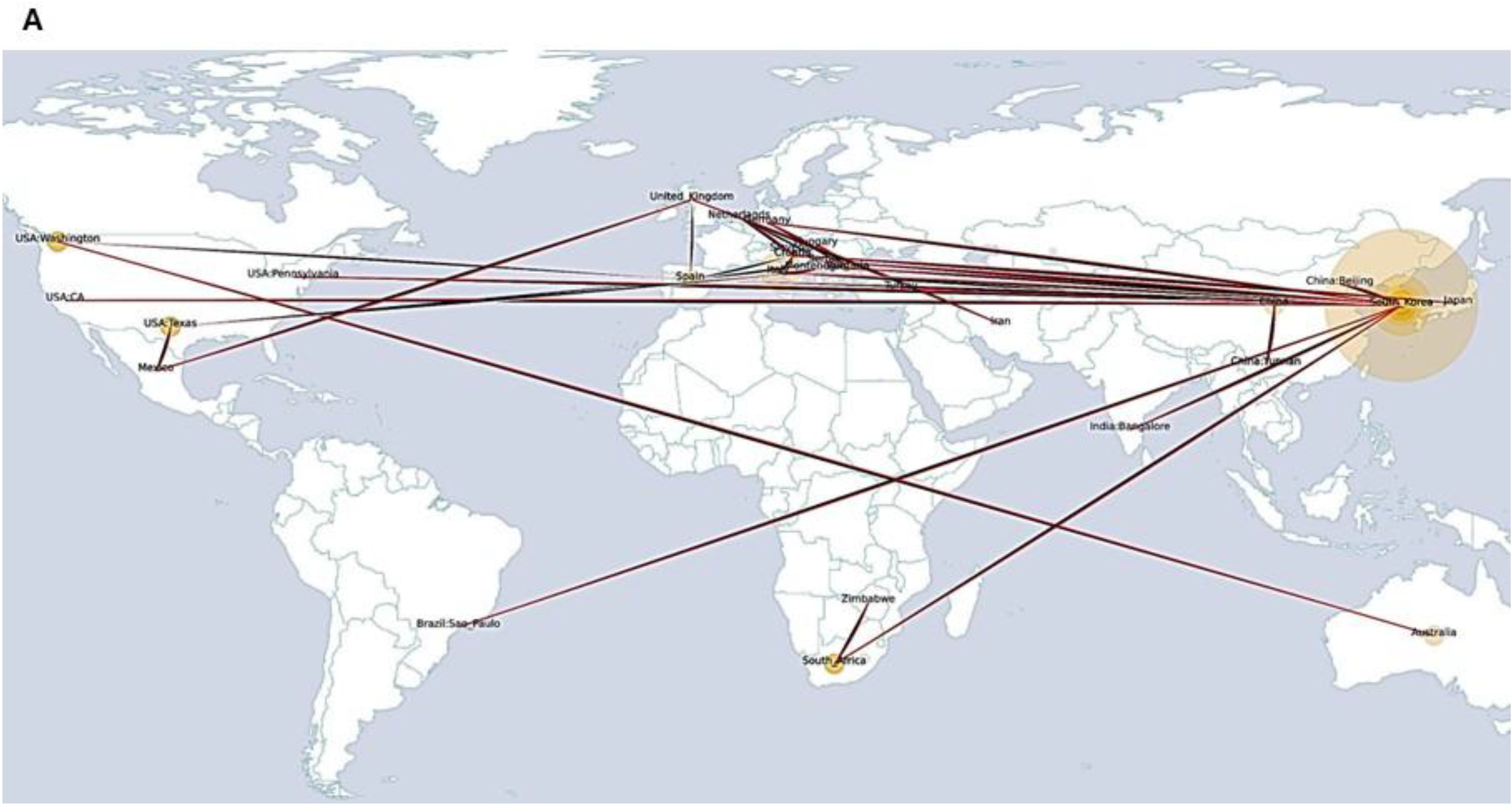
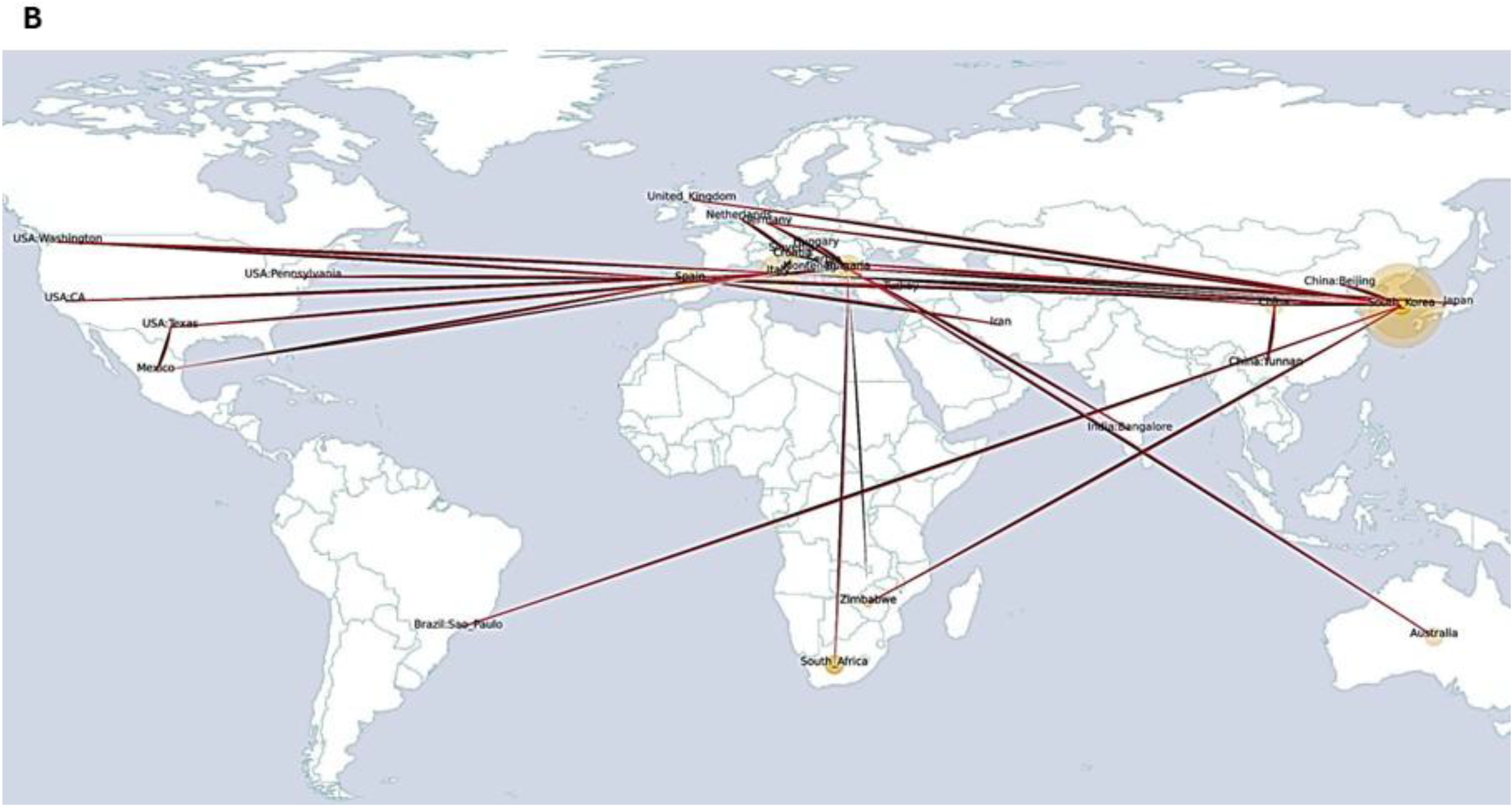
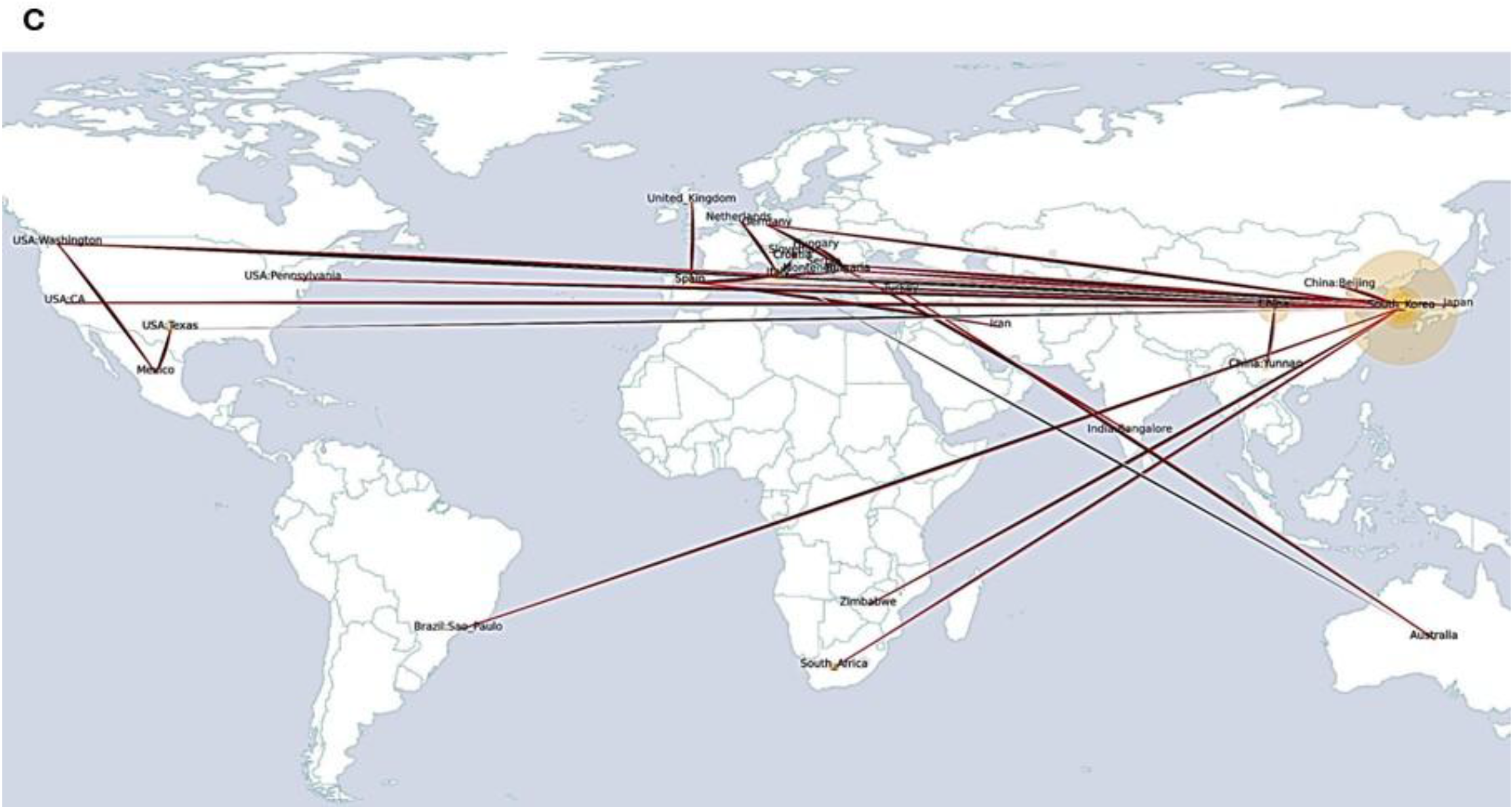
Spatial migration dynamics of TSWV inferred from whole genome sequences of A) Segment L, B) Segment M, and C) Segment S. The large internal yellow node indicates an expected ancestral state that corresponds to the geographic origin of TSWV, and smaller nodes indicate other localities were the TSWV has migrated. Red lines represent inferred dispersal routes indicating the locations of origin and destination.

In addition, we inferred the demographic history of TSWV simultaneously with the phylogeographic analysis, using the sequence data of the three genomic segments in separate analyses because each segment may follow different demographic courses. The Bayesian Skyline Plot (BSP) results showed similar patterns for the three genomic segments with slight differences. The BSP of the three segments showed that, after the initial emergence, a period of stability occurred as constant growth. Then, it was followed by two different periods of expansion and a final short phase of decline (Figure 4). An initial expansion occurred around 1900 for segments L and S, but for segment M, it came a little later (about 1940), and the second expansion occurred between 1990s-2000s for all segments. At the end of the record, we can see a slight contraction.

**Figure 4.**
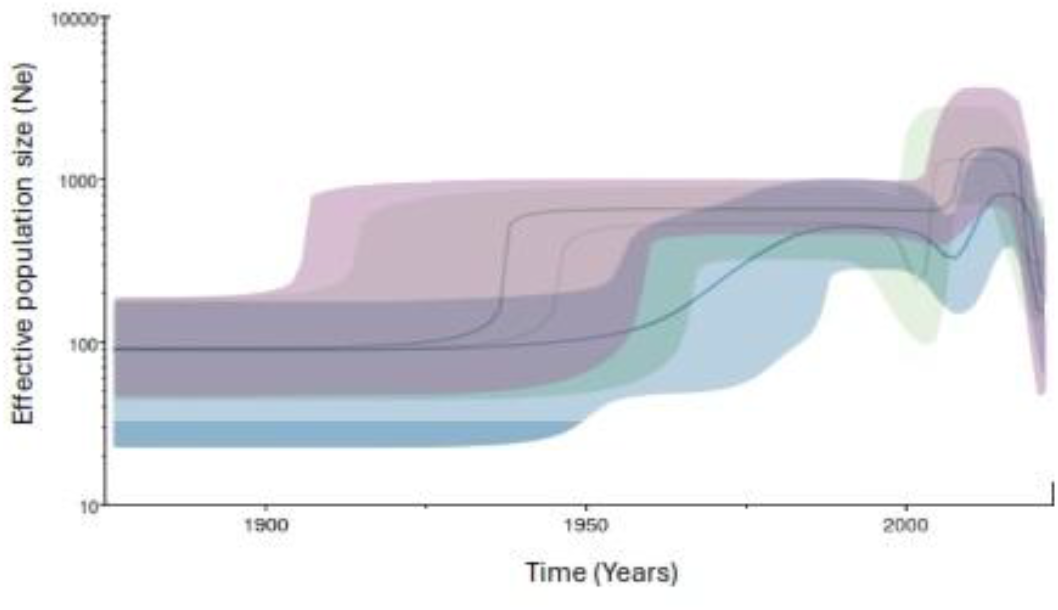
The demographic history of three TSWV genome segments was inferred via the Bayesian skyline plot (BSP) with a relaxed clock prior to all genomic segment data. The shading represents the 95% highest posterior density (HPD) of the product of generation time (τ) and effective population size (Ne). The solid line tracks the inferred median value. Color significance of the curves: Segment L: blue; Segment M: green; Segment S: purple.

Integrating the results of phylogeography with those of demographic dynamics, we can observe that the first significant migration of TSWV to Europe occurred around 1900, with the first strains arriving in the countries mentioned above about 1910. This date coincides with the first population expansion of TSWV observed in BSP. According to BSP, the second major expansion of the TSWV population occurred around 2000. By this date, the virus had already been distributed throughout almost all the localities in this investigation.

### Identification of possible ancestral virus that originated TSWV

Through the implementation of several phylogenetic analyses based on a single gene and full genome sequences, Butković and collaborators (2021) estimated that the relatives of TSWV are a group of tospoviruses, namely ANSV, CSNV, GRSV, TCSV, and ZLCV, (see Methods for virus full names). We further investigated to define which of these viruses would be the closest ancestor of TSWV by estimating genetic similarity and possible events of recombination. For this, we first reconstructed the ancestral sequences of five proteins encoded in the TSWV genome. The ancestral protein sequences inferred with FireProt-ASR do not differ substantially from current protein sequences (97-99% similar). This similarity is mainly because the current taxonomic classification of Tospovirus isolates designates viruses with ≥90% nucleotide similarity in the nucleocapsid gene to belong to the same species (Tsompana & Moyer, 2008). Since the present and ancestral nucleocapsid sequences are similar to each other, so are the respective sequences of the other proteins encoded in the TSWV genome. We performed a sliding windows analysis to determine the genetic similarity between TSWV and other tospoviruses. Results are concordant since both global and local similarities are higher when comparing TSWV with ANSV or CSNV than with other tospoviruses (see Table S3 and Table S4). Regarding global similarity, three proteins (RdRp, NSm, and glycoprotein precursor) show a higher similarity with ANSV, and two proteins (NSs and glycoprotein precursor) with CSNV. On the contrary, the N protein (nucleoprotein) deviates from this pattern because it presents higher similarity with GRSV and TCSV. When comparing the local genetic similarity of TSWV and related viruses, we found a higher number of similar local regions with ANSV or CSNV than other viruses, indicating that these viruses are more like TSWV; therefore, either one may have been the ancestral virus. To confirm this observation of ancestry, different phylogenetic reconstructions were made for each of the proteins. Figure S3 shows the results. ANSV constitutes the closest branch to TSWV in the tree constructed with RdRP protein sequences; however, both viruses formed distinct branches in the trees inferred with the other protein data. Although ANSV is closely related to TSWV in these trees, other viruses, such as TCSV and GRSV, are also well related.

From the sliding window analysis reported above, we can infer that several recombination events would have occurred between these viruses because the curves are thoroughly exchanged (mainly in the RdRp protein). We used the Proportion test to protein sequence data to test whether these viruses underwent recombination with each other. This tool is helpful to identify possible recombination events among protein sequences (Samson, 2022). The results show that TSWV would have undergone recombination with ANSV and other viruses to a similar extent (Table S5). That is, there is no clear predominance of ANSV in recombining with TSWV over the other viruses analyzed in this work.

## Discussion

Studying the evolution of plant viruses is crucial because it allows us to understand the dynamics of viral infections and their adaptation to new environments and plant hosts. This knowledge is essential for developing effective treatments and control strategies and the epidemiological surveillance of emerging viral diseases. In this study, we analyzed the evolutionary origin of one of the most devastating viruses for agricultural production, the tomato spotted wilt virus. Mainly, we focused on answering basic questions such as: what is the closest ancestral virus from which TSWV may have diverged? When did this virus happen, infecting various plants and crops? Where would have been the most likely place of its appearance? How has the TSWV population size changed over time? The abundance of genomic sequences available in databases and the recent generation of robust Bayesian inference programs now allow us to answer these questions.

The phylogenetic work of Butković and collaborators (2021) helped us narrow down the large number of possible viral ancestors of TSWV. Using sequence similarity, recombination analyses, and phylogenetic reconstruction, we deduced that two viruses -ANSV and CSNV-are the most likely candidates to be the ancestor of TSWV. Both viruses are classified in the TSWV genus, the Orthotospovirus, in the family Tospoviridae. ANSV might be the closest of the two since it shows more considerable global and local sequence similarity with TSWV. We confirmed this appreciation with phylogenetic reconstructions using protein sequences (Figure S3). The phylogeny indicates that ANSV is the closest branch to TSWV in the tree constructed with RdRP protein sequences; however, both viruses formed distinct branches in the trees inferred with the other protein data. Surprisingly, the relative abundance of recombination of TSWV with ANSV or CSNV is not higher than with other tospoviruses, suggesting that the ancestral TSWV was probably the original viral template that -through recombination and segment reassortment-gave rise to present TSWV, and perhaps ANSV, CSNV, or other viruses as well.

It is usually problematic to infer the divergence time of a species when nucleotide sequences do not naturally contain sufficient temporal information. We exhaustively searched for signals of temporality (commonly called the ‘clocklikeness’) in our sequences. Two robust methods clearly showed the clock-like character of the sequences of the three TSWV genomic segments. The BETS approach strongly supported heterochronous over isochronous models for all three sequence datasets. However, segment S showed a lower BF value, but still decisive, compared to the values of the other two segments, probably because the sequence length of segment S is shorter (∼2 to 3 times than the sequence of segments M and L). It is well-recognized that shorter sequences tend to have a lower temporal signal because they contain fewer genetic changes over time, leading to weaker correlations between genetic divergence and sampling time (Duchene et al., 2020). Further, analyses with the Date-Randomization Test showed that the TSWV sequences satisfied the most stringent criterion that confirms no overlap between the 95% credible interval of the actual rate estimate and any of those from the date-randomized data sets. These results indicate that the data contains enough temporal signal to calculate time-related issues. The Bayesian analyses revealed that the most recent common ancestor of TSWV diverged between 1768 and 1843. The 95% credibility interval width of the tree height parameter used for the tMRCA calculation is narrow for the Bayesian inferences performed with the three sequence datasets. This finding indicates a higher degree of confidence about the parameter being estimated, reflecting less uncertainty about the parameter’s actual value (Hespanhol et al., 2018). Among all three 95% credibility interval width values, the one for segment L is the smallest, perhaps because this dataset contains a stronger temporal signal. Therefore, we preferably take the year 1768 as the most probable date of the divergence of TSWV from its ancestor.

The estimation of the evolutionary rate, the tMRCA, and other evolutive parameters (see below) varies according to the sequence of the data set used for the calculations. This variation may be due to the temporal signal strength each sequence dataset contains, as indicated above, but also to the very nature of the tospovirus genome. Each genomic segment could follow a particular evolution mainly because TSWV unequally packages its genomic segments in the capsid (Michalakis & Blanc, 2024). The TSWV does not have a mechanism that ensures the even incorporation of a complete set of RNA segments into the same viral particle. Instead, genomic segments may be packaged in different viral particles that are independently transmitted (Michalakis & Blanc, 2020). Although TSWV is considered a monopartite virus (the different genome segments are packed in a single capsid), it adopts a multipartite lifestyle within the plant (Yvon et al., 2023). These differences in genomic content generate a sizeable genetic heterogeneity that, in this work, is reflected in marked differences in evolutionary rates and other evolutionary parameters.

Having said this, it is perhaps inappropriate to compare the phylogenies of the segments L, M, and S because they could have followed different evolutions, even though these segments belong to the same viral species. However, this is indeed what we found when we compared the phylogeny of each segment with that of the other. The disagreements in the tree topologies and branch lengths are significant (measured by the RF indices and Euclidean distances). These differences may also be due to the presence of traces of recombination in the data set. Although we have tried to reduce the presence of recombinants as much as possible, cryptic recombinants may still exist in the datasets used in this work. Further analyses must be done using larger sequence datasets of non-recombinant isolates to rule out the influence of recombination. Alternatively, analytical methods that take recombinants into account, such as phylogenetic networks, can be used to achieve a more comprehensive understanding of the TSWV phylogeny.

Phylogeographic dynamics analysis indicates that the origin of TSWV is South Korea. The phylogenetic trees reconstructed with the three datasets are rooted in South Korea, where the virus has spread worldwide. According to the BF analysis, the dispersion pathways appear robust because most pathways have high BF values that can be considered positive or strong, according to Kass & Raftery (1995). Although the geographic origin seems clear and well established, the spread to other localities has followed an arrangement that slightly differs according to the set of sequences used for the analysis. Very early, the virus may have reached Europe from South Korea and then North America and Africa (see Figure 3). Although we have 28 geographic locations in the TSWV sample, a broader phylogeographic study should be carried out to include many locations to have a more complete picture of the spread of the virus worldwide.

Demographic analysis indicates that TSWV experienced two major population increases. The first population expansion can be explained by the migration of the virus to Europe and subsequent localities that may have opened new horizons of infection. The availability of new plant hosts may have increased the population significantly. In this sense, the intensification of agricultural production, the use of monocultures, and the expansion of new lands with the consequent land conversion may have triggered the emergence of TSWV (Rojas & Gilbertson, 2008). As said, TSWV has an extensive host range at present, with over 1000 plant species infected from more than 90 different plant families (Kormelink et al., 2011). Likewise, the possibility of finding new vector species in different localities may have prompted population growth. TSWV is transmitted by various species from the order Thysanoptera (Hogenhout et al., 2008). It is well known that the proliferation of tospoviruses is preceded by increases in the distribution and population of insect vectors (Rojas & Gilbertson, 2008). Early reports from the 1930s indicated that *Frankliniella occidentalis* was the vector of TSWV in the USA, however later registers informed that *F. fusca* and *Thrips palmi, T. tabaci*, and *T. setosus* have also been found to transmit TSWV at different efficiencies (Chatzivassiliou et al., 2000; Inoue et al., 2007). In the late 1980s, there was a resurgence of TSWV in the USA and Western Europe (Jones, 2005). This new outbreak and increase in disease caused by TSWV have been attributed to the spread of *F. occidentalis* from the Western USA to other areas of the world (Marchoux et al., 1991). A few years later, new *Frankliniella* species were reported as vectors of TSWV. In 1995, *F. schultzei* and *F. intonsa* were confirmed as TSWV vectors; in 1998, *F. bispinosa* was shown to transmit TSWV. In 2004, *F. intonsa, F. occidentalis, T. tabaci* and *T. setosus* were confirmed as vectors too (Jones, 2005). This outburst coincides with the second-largest population increase of TSWV, according to the plots outlined with the results of the Bayesian Skyline analysis. However, we cannot rule out other causes that may have influenced the excessive growth of TSWV around the year 2000.

Our study provides new insights into the evolutionary origin and the demographic history of TSWV. In particular, we estimated the most probable viral ancestor of TSWV and the approximate date when the divergence from this ancestor might have happened. We also found the geographic origin of this virus and the multiple migration pathways that TSWV used to spread worldwide. These results provide critical epidemiological data to design control strategies for this virus or other similar species. Although this work is the first attempt to understand the virus’s origin, phylogeography, and demography, further analyses that consider recombination and reassortment are required. Similarly, to get a more detailed understanding of the TSWV geographic spread, future work should include an extensive collection of isolates sampled from many countries and localities.

## Methods

### Nucleotide sequences, alignment, and recombination detection

Whole genome sequences of TSWV were downloaded from the NCBI Virus database in August 2024. The raw dataset was composed of 800 sequences corresponding to each genome segment. We organized the data to have sequences of the three segments from the same TSWV isolates. The sequences were aligned with MAFFT v. 7 (Katoh et al., 2019), producing three different alignments (for segments L, M, and S) of 153 sequences each. Sequences were then examined with the RDP v.5.58 package (Martin et al., 2020) to detect evidence of recombination. We run seven different methods (RDP, bootscan, maxchi, chimaera, 3seq, geneconv, and siscan) with Bonferroni correction and set the *p*-value to 0.05 to detect recombination. All sequences showing recombination signals were removed from the datasets for a streamlined phylogenetics-based molecular evolution analysis. Only the sequences detected as recombinants by five or more methods were considered authentic recombinants and eliminated from the dataset. The removal of recombinants resulted in a reduced dataset of 136 sequences for each of the genomic segments, which is the dataset that was used for all successive analyses. We paid attention to keeping information about the sampling year in each sequence (heterochronous sequences) to be able to perform time-related analyses. Table S1 summarizes the information on the TSWVs, including database accession numbers of genomic sequences.

We assessed the substitution saturation of TSWV sequence alignments with the software DAMBE (Xia & Xie, 2001) to verify the sequences’ suitability for performing phylogenetic analysis. We calculated the *Iss* (index of substitution saturation) for all sites and for resolved sites, and statistically, we compared them with the critical value of the index of substitution saturation (*Iss*.*c*).

### Analysis of temporal signal

The isolation sampling window of the TSWV dataset analyzed in this study is 35 years. We inquired whether this time interval was sufficient to accumulate temporal information to perform temporal-related evolutionary analyses. We tested the temporal signal that the TSWV sequences may have using two different methods: 1) performing the Bayesian Evaluation of Temporal Signal (BETS), which is aimed at estimating the ratio of the marginal likelihoods of two competing models, one model calculated using the actual sampling times (i.e., heterochronous) and another model in which the sampling times were - arbitrarily-adjusted to have the exact sampling times (i.e. isochronous, Duchene et al., 2020); 2) performing the date-randomization test which involves generating multiple data sets with randomly altered sampling times and compared to actual data sets (Duffy & Holmes, 2009). For the first above-mentioned analysis, we quantified the statistical support of the two competing models (i.e., heterochronous vs. isochronous) by calculating the marginal likelihood using the path sampling (PS) method implemented in the package BEAST 2.7.7 (Bouckaert et al., 2019). For PS, we adjusted the number of steps to 100 and used a chain length equal to 200000 (we experimentally determined the chain length for consistency by running many PS calculations). The marginal likelihood results helped estimate the log Bayes Factor (logBF). The Bayes factor is the ratio of marginal likelihoods, expressed in this way: logBF=log(P(*Y*|M_model_A_))-log(P(*Y*|M_model_B_)). A logBF of 0-2 means weak evidence, 2-6 denotes positive evidence and 6-10 indicates strong support for one model over the other (Kass & Raftery, 1995). For the second analysis, we calculated the clock rate of the actual dataset and the “randomized” dataset using BEAST 1.10.4 (Suchard et al., 2018). We run one analysis for the real dataset and ten different “randomized” datasets, setting the parameters to the strict clock with a constant size tree prior. The Markov chain Monte Carlo (MCMC) analyses were run for 30 or 40 million with sampling every 1000 steps. Results were analyzed in Tracer 1.7.1 (Rambaut et al., 2018) to ensure all parameters reached equal or over 200 effective sample sizes. Then, we compared the mean and the 95% HPD intervals of clock rate to confirm that the mean rate estimate of the sample with correct dates does not overlap with any of the 95% credible intervals of ten samples with randomized dates.

### Temporal evolution of TSWV

To estimate the evolutionary rate and the time to the most recent common ancestor (tMRCA) of TSWV, we first examined which clock model and tree prior best fit the data using BEAST 2.7.7. We analyzed two clock models (strict and relaxed clock) and three demographic models (constant size, exponential growth, and Bayesian skyline) for the datasets corresponding to segments L, M, and S in independent analyses. We calculated the marginal likelihood estimate (MLE) using PS for model selection. For PS, we employed a chain length of 200000, 100 steps, and a burnin of 50%. We calculated the BF as stated above using the PS estimates for each genomic segment and the respective combination of clock and tree models to obtain the results in Table 1. All Bayesian phylogenetic analyses were set up in BEAST to GTR, G4+I, with empirical frequencies for sites and ORC algorithm (Douglas et al., 2021) for the relaxed clock to reach faster convergence. The best-fit model of nucleotide substitution was calculated using the ModelFinder Plus (MFP) tool in IQTree 2.3.6 (Minh et al., 2020). Coincidentally, this model was the same for each genome segment dataset. The analyses were run for 30 to 100 million steps across MCMC simulations and sampled every 1000 or 4000 steps, depending on the dataset. We also included control analysis without sequence data to avoid taking the results obtained from priors as valid. We discarded 10% of the samples as burnin. Only values of ESS ≥200 were considered acceptable (i.e., the analysis reached convergence) when log files were analyzed in Tracer 1.7.1 (Rambaut et al., 2018). To obtain the best tree from a tree file, we searched for the maximum clade credibility tree using the TreeAnnotator program (Heled & Bouckaert, 2013) and then visualized it with FigTree (Rambaut, 2018).

### Demographic and discrete phylogeography analyses

The demographic history of TSWV was estimated using the Bayesian skyline model implemented on BEAST 2.7.7. We ran analyses for the sequences of the three genome segments in combination with the best clock model deciphered with BF (relaxed in this case) using the parameter settings above described. The Tracer program was used for demographic reconstruction to obtain the Bayesian Skyline Plot (BSP). For this, we used the complete set of tree data inferred with BEAST and set it up to 2023, which is the date of the youngest tip.

The geographical origin and the inter-regional (including inter-continental) spread of TSWV were estimated from segments L, M, and S sequence samples via Bayesian discrete phylogeography using the BEAST 2.7.7 package. The sampling locations were considered discrete traits, and geographic migration is inferred based on the evolutionary tree. We employed phylogeographic analysis together with the Skyline model, as this tree prior turned out to be the best for TSWV samples. Phylogeography with Skyline and molecular clock helps to infer both spatial and temporal transmission dynamics of the virus, which is feasible to perform in the Beast 2.7.7. The different priors of tree and clock models (Skyline and ORC) were set up as described above. We used a symmetric matrix for the rate of change between two states that, in this case, are transition rates between all pairs of sampled locations. In the symmetric model, both forward and backward rates are estimated from the sample. The TSWV dataset has 28 different locations, and large countries (e.g., USA, China, India) are split into internal locations (California, Texas, Pennsylvania, Yunnan, etc.). Calculating rates is usually difficult since genomic data is unlikely to contain enough information about all possible transitions. Therefore, we computed the BF to support the posterior rate indicator expectation. BF adds robustness to migration rate estimation in the geographic spread of TSWV. We ran Beast analyses with 40 or 60 million MCMC chain lengths. The first 10% of the samples from the MCMC chain were discarded as burn-in, and the remaining trees were summarized as a maximum clade credibility tree using TreeAnnotator 2.7.7. The Spread utility (Nahata et al., 2022) was helpful in visualizing results in a world map and calculating the BF as the transition rate support.

### Ancestral sequence estimation and calculation of genetic similarity between tospoviruses

To identify the ancestral virus from which the current TSWV could have been derived, we downloaded the protein sequences encoded in the three genomic segments of the 136 TSWV isolates. These are five proteins, namely the nucleoprotein (N) and the protein NS_s_ (from segment S), the NS_m_ and the precursor of glycoproteins G_N_ and G_C_ (from segment M), and the RNA-dependent RNA polymerase (RdRp, from segment L). We reconstructed the ancestral amino acid sequences of these five proteins in separate analyses using the web server FireProt-ASR (https://loschmidt.chemi.muni.cz/fireprotasr/) that allows the automated estimation of ancestral protein sequences (Musil et al., 2020). The phylogenetic tree generated by this program enables the selection of sequences from ancestral nodes. We chose the five sequences closest to the root. We also obtained the respective protein sequences from other TSWV-related tospoviruses, that is to say, from Alstroemeria necrotic streak virus (ANSV), Chrysanthemum stem necrosis virus (CSNV), Groundnut ringspot virus (GRSV), Tomato chlorotic spot virus (TCSV) and Zucchini lethal chlorosis virus (ZLCV). Protein sequences from two isolates of each relative virus were downloaded from the database and merged into a single fasta file with the five ancestral protein sequences of TSWV. We then aligned the protein sequence datasets with the MAFFT Multiple alignment program. Each protein alignment was used for a sliding window analysis to estimate the genetic similarity between TSWV and the TSWV-related tospoviruses to search for possible ancestors and to detect recombination among them. For this purpose, we employed the SimPlot++ program (Samson et al., 2022), setting the window length to 100 aa for proteins N, NS_s_, and NS_m_ and 200 aa for RdRp and glycoprotein precursors because the latter are larger proteins. The strip gap was set up to 33%, and the JTT92 distance model was used for all calculations.

Maximum likelihood (ML) phylogenetic trees were inferred in IQTree v. 2.3.6 (Minh et al., 2020) using sequences of the five proteins encoded by the three viral genomic RNAs. We included current and ancestral sequences in the phylogenetic reconstruction as described above. We searched for the best nucleotide substitution model with the -m TEST option, and branch support was calculated with 1000 replicates of ultrafast bootstrap (UFBoot). The UFBoot trees were optimized with nearest neighbor interchange to reduce the risk of overestimating branch statistics due to possible model violations. The branch support was also assessed employing the Shimodaira–Hasegawa-like approximate likelihood ratio test. The ML trees were graphed with Figtree v.1.4.4 (Rambaut, 2018).

## Supporting information

Table S1. Tomato spotted wilt virus isolates used in this study Table S2. Putative major recombination sites found in genome segments of TSWV. Table S

Figure S1. Side-by-side comparison of phylogenetic trees reconstructed with whole genome sequences of TSWV segments L, M, and S. The blue-to-yellow co

## Acknowledgements

We thank Yachay Tech University, Ecuador, for partial funding through internal grant PGI24-02 to J.A.C. and the Phage Therapy Group. We also thank Ms. Helen Guigues for her valuable contribution to data processing and analyses.

## Data availability

Sequence data and geographic coordinates used in this work are described in Table S1, Supplementary Material.

## Competing interests

None declared.

## Notes

### Competing Interest Statement

The authors have declared no competing interest.

