## Supplementary material for "Evolutionary origin and demographic trajectory of tomato spotted wilt virus (*Orthotospovirus tomatomaculae*)": Table S1. Tomato spotted wilt virus isolates used in this study Table S2. Putative major recombination sites found in genome segments of TSWV. Table S

| Isolate name | Accession number | | | Year of sampling | Place of sampling | Latitude | Longitude | Plant Host |
|---|---|---|---|---|---|---|---|---|
| | Segment L | Segment M | Segment S | | | | | |
| 100/23 | PP968744.1 | PP968745.1 | PP968746.1 | 2023 | Montenegro | 42.720 | 19.380 | Tomato |
| 102SEC22S | OQ534348.1 | OQ534347.1 | OQ534346.1 | 2022 | Croatia | 45.149 | 15.195 | Cucumber |
| 104DOT22S | OQ269472.1 | OQ534350.1 | OQ534349.1 | 2022 | Croatia | 45.149 | 15.195 | Tomato |
| 105DOT22S | OQ507123.1 | OQ507122.1 | OQ507121.1 | 2022 | Croatia | 45.149 | 15.195 | Tomato |
| 106DOT22S | OQ534342.1 | OQ534341.1 | OQ534340.1 | 2022 | Croatia | 45.149 | 15.195 | Tomato |
| 107DOT22S | OQ507126.1 | OQ507125.1 | OQ507124.1 | 2022 | Croatia | 45.149 | 15.195 | Tomato |
| 108DOT22S | OQ507129.1 | OQ507128.1 | OQ507127.1 | 2022 | Croatia | 45.149 | 15.195 | Tomato |
| 109DOP22S | OQ507132.1 | OQ507131.1 | OQ507130.1 | 2022 | Croatia | 45.149 | 15.195 | Pepper |
| 253SEP23S | PP943378.1 | PP943379.1 | PP943380.1 | 2023 | Croatia | 45.149 | 15.195 | Pepper cv. Prince F1 |
| 254SEP23S | PP943381.1 | PP943382.1 | PP943383.1 | 2023 | Croatia | 45.149 | 15.195 | Pepper cv. Prince F1 |
| 255_256SEP23S | PP943384.1 | PP943385.1 | PP943386.1 | 2023 | Croatia | 45.149 | 15.195 | Pepper cv. Dragoney |
| 52STT21S | OP373186.1 | OP373185.1 | OP373184.1 | 2021 | Croatia | 45.149 | 15.195 | Tomato |
| 71SET22S | OQ534345.1 | OQ534344.1 | OQ534343.1 | 2022 | Croatia | 45.149 | 15.195 | Tomato |
| 85DOPt22S | OQ507117.1 | OQ507116.1 | OQ507115.1 | 2022 | Croatia | 45.149 | 15.195 | Potato |
| 86DOW22S | OQ507120.1 | OQ507119.1 | OQ507118.1 | 2022 | Croatia | 45.149 | 15.195 | *Amaranthus blitum* |
| 98/23 | PP968741.1 | PP968742.1 | PP968743.1 | 2023 | Montenegro | 42.720 | 19.380 | Tomato cultivar Pink Gusto |
| Ag_SA1 | OP921761.1 | OP921762.1 | OP921763.1 | 2021 | South Africa | -30.387 | 22.894 | *Agapanthus praecox* |
| BasC | MH745370.1 | KU179514.1 | KU179515.1 | 2014 | USA: Washington | 47.763 | -120.740 | *Ocimum basilicum* |
| Bidens | MF805766.1 | MF805765.1 | MF805764.1 | 2016 | China: Yunnan | 25.142 | 10.269 | Bidens |
| Blackberry_lily-kr1 | MT899477.1 | MT921847.1 | MT921846.1 | 2020 | South Korea | 35.958 | 12.778 | *Iris domestica* |
| BR | MF159038.1 | MF159050.1 | MF159062.1 | 2016 | South Korea | 35.958 | 12.778 | Pepper |
| CY | MF159039.1 | MF159051.1 | MF159063.1 | 2016 | South Korea | 35.958 | 12.778 | Pepper |
| CY-G1-2 | MT643234.1 | MT842839.1 | MN854655.1 | 2018 | South Korea | 35.958 | 12.778 | *Lycium chinense* |
| Daegok_Jinju | OP595386.1 | OP595390.1 | OP595394.1 | 2022 | South Korea | 35.958 | 12.778 | Pepper |
| DG | LC790665.1 | LC790666.1 | LC790667.1 | 2023 | South Korea | 35.958 | 12.778 | *Arctium lappa* |
| DSMZ_PV-0182 | MZ202328.1 | MZ202329.1 | MZ202330.1 | 1988 | Bulgaria | 42.779 | 25.450 | Tobacco |
| DSMZ_PV-0389 | MT723985.1 | MT723986.1 | MT723987.1 | 1994 | Brazil: Sao Paulo | -23.549 | -46.639 | *Nicotiana rustica* |
| DSMZ_PV-0393 | MW854271.1 | MW854272.1 | MW854273.1 | 1993 | Bulgaria | 42.779 | 25.450 | Tobacco |
| DSMZ_PV-1175 | ON924222.1 | ON924223.1 | ON924224.1 | 2015 | Hungary | 47.199 | 19.507 | Pepper |
| DSMZ_PV-1369 | OQ102068.1 | OQ102069.1 | OQ102070.1 | 2022 | Netherlands | 52.170 | 52.870 | *Callistephus* |
| DSMZ_PV-1412 | OR762544.1 | OR762545.1 | OR762546.1 | 2023 | Serbia | 44.016 | 21.006 | Pepper |
| DSMZ_PV-1413 | OR762547.1 | OR762548.1 | OR762549.1 | 2023 | Turkey | 39.059 | 35.250 | Tomato |
| DSMZ_PV-0204 | MT682292.1 | MT682293.1 | MT682294.1 | 1988 | Germany | 51.250 | 10.460 | Impatiens New-Guinea hybrid |
| DSMZ_PV-1265 | ON924228.1 | ON924229.1 | ON924230.1 | 2019 | Germany | 51.250 | 10.460 | Pepper |
| DY | MF159040.1 | MF159052.1 | MF159064.1 | 2016 | South Korea | 35.958 | 12.778 | Pepper |
| eggplant | OM154968.1 | OM154967.1 | OM154966.1 | 2021 | China: Beijing | 39.910 | 116.400 | Eggplant |
| FC-19 | LC652640.1 | LC652641.1 | LC652642.1 | 2019 | Japan | 36.330 | 138.268 | *Cucumis sativus* |
| FERA_18-1.23 | OM112200.1 | OM112201.1 | OM112202.1 | 2018 | United Kingdom | 55.550 | -3.450 | Tobacco |
| FG-tomSRB | OR166268.1 | OR166267.1 | OR166266.1 | 2022 | Italy | 41.980 | 12.540 | Tomato cv. Docet |
| G | MF159041.1 | MF159053.1 | MF159065.1 | 2016 | South Korea | 35.958 | 12.778 | Pepper |
| GC | MF159042.1 | MF159054.1 | MF159066.1 | 2016 | South Korea | 35.958 | 12.778 | Pepper |
| Ger | LC603056.1 | LC535227.1 | LC535228.1 | 2019 | South Korea | 35.958 | 12.778 | *Gerbera hybrida* |
| Geumsan_Jinju | OP595384.1 | OP595388.1 | OP595392.1 | 2022 | South Korea | 35.958 | 12.778 | Pepper |

| | | | | | | | | |
|---|---|---|---|---|---|---|---|---|
| Goesan | OM022891.1 | OM022883.1 | OM022881.1 | 2021 | South Korea | 35.958 | 12.778 | Pepper |
| GS | MF159043.1 | MF159055.1 | MF159067.1 | 2016 | South Korea | 35.958 | 12.778 | Pepper |
| Gumi | MW048592.1 | MW048591.1 | MW048590.1 | 2019 | South Korea | 35.958 | 12.778 | *Gerbera jamesonii* |
| Hahoe_Andong | OM022887.1 | OM022882.1 | OM022877.1 | 2021 | South Korea | 35.958 | 12.778 | Pepper |
| Henan | MT799177.1 | MT799178.1 | MT799179.1 | 2020 | China | 36.040 | 104.220 | Pepper |
| HLJ-1 | MG878873.1 | MG878874.1 | MG878875.1 | 2017 | China | 36.040 | 104.220 | Tomato |
| HS | MF159044.1 | MF159056.1 | MF159068.1 | 2016 | South Korea | 35.958 | 12.778 | Pepper |
| I | MF159045.1 | MF159057.1 | MF159069.1 | 2016 | South Korea | 35.958 | 12.778 | Pepper |
| IRP4 | PP780924.1 | PP780925.1 | PP780926.1 | 2023 | Iran | 32.450 | 53.670 | Pepper |
| Jeonju | OM022888.1 | OM022884.1 | OM022879.1 | 2021 | South Korea | 35.958 | 12.778 | Pepper |
| JJ | MF159046.1 | KY021438.1 | KY021439.1 | 2016 | South Korea | 35.958 | 12.778 | Pepper |
| K | MF159047.1 | MF159059.1 | MF159071.1 | 2016 | South Korea | 35.958 | 12.778 | Pepper |
| Ka-To | MK977648.1 | MK977649.1 | MK977650.1 | 2019 | India: Bangalore | 12.970 | 77.590 | Tomato |
| Kunming | OM902668.1 | OM902667.1 | OM902666.1 | 2014 | China: Yunnan | 25.142 | 10.269 | Tomato |
| LK-1 | KY250488.1 | KY250489.1 | KY250490.1 | 2015 | South Africa | -30.387 | 22.894 | *Amaranthus thunbergii* |
| LL-N.05 | KP008128.1 | FM163373.1 | KP008129.1 | 2005 | Spain | 40.730 | -3.750 | Tomato |
| LN-HJL | MT241883.1 | MT241884.1 | MT241885.1 | 2019 | China | 36.040 | 104.220 | *Tropaeolum majus L.* |
| Mexico | OQ718495.1 | OQ718496.1 | OQ718497.1 | 2016 | Mexico | 23.850 | -102.590 | Tomato |
| Micheon_Jinju | OP595385.1 | OP595389.1 | OP595393.1 | 2022 | South Korea | 35.958 | 12.778 | Pepper |
| MR-01 | MG593197.1 | MG593198.1 | MG593199.1 | 2015 | USA: CA | 36.880 | -119.390 | *Cichorium intybus* |
| Mum-A5 | MG602671.1 | MG602672.1 | MG602673.1 | 2014 | Zimbabwe | -18.950 | 29.160 | *Chrysanthemum* |
| Non-RB | PP632110.1 | PP632111.1 | PP632112.1 | 2023 | USA: Texas | 32.060 | -99.890 | Pepper |
| NS-AG28 | MT643235.1 | MT842840.1 | MN854653.1 | 2019 | South Korea | 35.958 | 12.778 | *Angelica gigas* |
| NS-BB20 | MT643236.1 | MT842841.1 | MN854654.1 | 2019 | South Korea | 35.958 | 12.778 | *Petasites japonicus* |
| p202/3WT | KJ575619.1 | HQ830188.1 | HQ830187.1 | 1999 | Italy | 41.980 | 12.540 | Pepper |
| PA01 | KT160280.1 | KT160281.1 | KT160282.1 | 2014 | USA: Pennsylvania | 41.250 | -77.180 | Pepper |
| Pap | MF326509.1 | MF326510.1 | MF326511.1 | 2016 | South Korea | 35.958 | 12.778 | Pepper |
| PepCal_10 | MH763621.1 | MH756624.1 | MG989673.1 | 2014 | Italy | 41.980 | 12.540 | Pepper |
| PepCal_12 | MK348941.1 | MH756625.1 | MG989674.1 | 2014 | Italy | 41.980 | 12.540 | Pepper |
| Pepper1_CY-CN | HM581937.1 | HM581938.1 | HM581939.1 | 2009 | South Korea | 35.958 | 12.778 | Pepper |
| Pepper2_CY-CN | HM581940.1 | HM581941.1 | HM581942.1 | 2009 | South Korea | 35.958 | 12.778 | Pepper |
| PLE20ST2 | OL471946.1 | OL471947.1 | OL471948.1 | 2020 | Slovenia | 46.170 | 14.990 | Tomato |
| PLE20ST3 | OL471949.1 | OL471950.1 | OL471951.1 | 2020 | Slovenia | 46.170 | 14.990 | Tomato |
| Pujol1TL3 | KP008130.1 | HM015520.1 | KP008131.1 | 2003 | Spain | 40.730 | -3.750 | Tomato |
| PVR | KP008132.1 | KP008133.1 | KP008134.1 | 2007 | Spain | 40.730 | -3.750 | Pepper |
| SDLY-1 | MN833242.1 | MN870629.1 | MN861978.1 | 2019 | China | 36.040 | 104.220 | Tobacco |
| SK-J | MZ404050.1 | MZ404051.1 | MZ404052.1 | 2020 | South Korea | 35.958 | 12.778 | Tobacco |
| Songcheon | OM022890.1 | OM022886.1 | OM022880.1 | 2021 | South Korea | 35.958 | 12.778 | Pepper |
| SS | LC685923.1 | LC685924.1 | LC685925.1 | 2021 | South Korea | 35.958 | 12.778 | *Lobelia erinus* cv. Sweet Springs |
| SW-TO2 | MT842843.1 | MT842842.1 | MT842844.1 | 2019 | South Korea | 35.958 | 12.778 | Tomato |
| T1012 | ON840010.1 | ON840011.1 | ON840012.1 | 2005 | Italy | 41.980 | 12.540 | Tomato |
| T1107 | ON840007.1 | ON840008.1 | ON840009.1 | 2021 | Italy | 41.980 | 12.540 | Tomato |
| Tomato_NJ-JN | HM581934.1 | HM581935.1 | HM581936.1 | 2008 | South Korea | 35.958 | 12.778 | Tomato |
| Tom-BL2 | PP622754.1 | PP622755.1 | PP622756.1 | 2022 | USA: Texas | 32.060 | -99.890 | Tomato |
| Tom-MX | PP622757.1 | PP632102.1 | PP632103.1 | 2022 | Mexico | 23.850 | -102.590 | Tomato |
| TSWV-10 | KC261962.1 | KC261963.1 | KC261964.1 | 2009 | South Korea | 35.958 | 12.778 | *Stellaria aquatica* |
| TSWV-12 | KC261965.1 | KC261966.1 | KC261967.1 | 2010 | South Korea | 35.958 | 12.778 | Lettuce |

| | | | | | | | | |
|---|---|---|---|---|---|---|---|---|
| TSWV-16 | KC261968.1 | KC261969.1 | KC261970.1 | 2010 | South Korea | 35.958 | 12.778 | Tomato |
| TSWV-17 | KC261971.1 | KC261972.1 | KC261973.1 | 2010 | South Korea | 35.958 | 12.778 | *Stellaria media* |
| TSWV-18 | KC261974.1 | KC261975.1 | KC261976.1 | 2010 | South Korea | 35.958 | 12.778 | *Chrysanthemum* |
| TSWV-4 | KC261947.1 | KC261948.1 | KC261949.1 | 2009 | South Korea | 35.958 | 12.778 | Pepper |
| TSWV-5 | KC261950.1 | KC261951.1 | KC261952.1 | 2009 | South Korea | 35.958 | 12.778 | *Stellaria aquatica* |
| TSWV-6 | KC261953.1 | KC261954.1 | KC261955.1 | 2009 | South Korea | 35.958 | 12.778 | *Stellaria media* |
| TSWV-7 | KC261956.1 | KC261957.1 | KC261958.1 | 2009 | South Korea | 35.958 | 12.778 | Pepper |
| TSWV-8 | KC261959.1 | KC261960.1 | KC261961.1 | 2009 | South Korea | 35.958 | 12.778 | *Lactuca indica* |
| TSWV-AS-P1 | LC742962.1 | LC742963.1 | LC742964.1 | 2022 | South Korea | 35.958 | 12.778 | Pepper |
| TSWV-BJFC-Cb | OM937132.1 | OM982913.1 | OM982912.1 | 2021 | China | 36.040 | 104.220 | *Coreopsis basalis* |
| TSWV-CY-LC | MN064722.1 | MN064723.1 | MN064724.1 | 2018 | South Korea | 35.958 | 12.778 | *Lycium chinense* |
| TSWV-HJ | LC273305.1 | LC273306.1 | LC273307.1 | 2016 | South Korea | 35.958 | 12.778 | *Humulus japonicus* |
| TSWV-LN | MK986669.1 | MK986670.1 | MK986671.1 | 2019 | China | 36.040 | 104.220 | Tomato |
| TSWV_Nto | OL471723.1 | OL471716.1 | OL471721.1 | 2019 | South Korea | 35.958 | 12.778 | Tobacco |
| TSWV-QLD1 | KT717691.1 | KT717692.1 | KT717693.1 | 2014 | Australia | -25.170 | 133.800 | *Capsicum sp.* cv. yolo wonder |
| TSWV-QLD2 | MG025802.1 | MG025803.1 | MG025804.1 | 2015 | Australia | -25.170 | 133.800 | Pepper cv. Warlock |
| TSWV-RP-GJ | OL742435.1 | OL742436.1 | OL742437.1 | 2021 | South Korea | 35.958 | 12.778 | Red Pepper |
| TSWV-YN | JF960237.1 | JF960236.1 | JF960235.1 | 2010 | China | 36.040 | 104.220 | Tomato |
| TUR20ST1 | OL471952.1 | OL471953.1 | OL471954.1 | 2020 | Slovenia | 46.170 | 14.990 | Tomato |
| TUR20ST2 | OL471955.1 | OL471956.1 | OL471957.1 | 2020 | Slovenia | 46.170 | 14.990 | Tomato |
| TUR20ST3 | OL471958.1 | OL471959.1 | OL471960.1 | 2020 | Slovenia | 46.170 | 14.990 | Tomato |
| TUR20SW | OL471961.1 | OL471962.1 | OL471963.1 | 2020 | Slovenia | 46.170 | 14.990 | Weed plants |
| Wa1 | MH745369.1 | KU179512.1 | KU179513.1 | 2013 | USA: Washington | 47.763 | -120.740 | Tomato |
| water_dropwort | OM154971.1 | OM154970.1 | OM154969.1 | 2021 | China: Beijing | 39.910 | 116.400 | *Oenanthe crocata* |
| WJ | MW293977.1 | MW293978.1 | MW293979.1 | 2020 | South Korea | 35.958 | 12.778 | Pepper |
| Yeongwol | OP595383.1 | OP595387.1 | OP595391.1 | 2022 | South Korea | 35.958 | 12.778 | Pepper |
| YI | MW293974.1 | MW293975.1 | MW293976.1 | 2020 | South Korea | 35.958 | 12.778 | Pepper |
| YKMiSFQ1 | MW404243.1 | MW404244.1 | MW404245.1 | 2018 | China: Yunnan | 25.142 | 10.269 | *Solanum betaceum* |
| YN5573 | MF590699.1 | MF590698.1 | KY495609.1 | 2016 | China:Yunnan | 25.142 | 10.269 | Pea |
| YN5574 | MF422030.1 | KY495606.1 | KY495610.1 | 2016 | China: Yunnan | 25.142 | 10.269 | *Codonopsis pilosula* |
| YN5575 | MF422031.1 | KY495607.1 | KY495611.1 | 2016 | China: Yunnan | 25.142 | 10.269 | Dahlia |
| YN5576 | MF590700.1 | KY495608.1 | KY495612.1 | 2016 | China:Yunnan | 25.142 | 10.269 | *Solanum lasiocarpum* |
| YN5577 | MF422032.1 | MF422033.1 | MF422034.1 | 2016 | China: Yunnan | 25.142 | 10.269 | *Tropaeolum majus* |
| YNgp | KM657122.1 | KM657119.1 | KM657116.1 | 2013 | China | 36.040 | 104.220 | Green Pepper |
| YNHHML | MN833247.1 | MN870632.1 | MN861981.1 | 2019 | China | 36.040 | 104.220 | Tobacco |
| YNHHMZ | MN870626.1 | MN870630.1 | MN861979.1 | 2019 | China | 36.040 | 104.220 | Tobacco |
| YNHS | MN365036.1 | MN365035.1 | MN365037.1 | 2018 | China | 36.040 | 104.220 | Peanut |
| YNKM-1 | MN865481.1 | MN870628.1 | MN861976.1 | 2019 | China | 36.040 | 104.220 | Tobacco |
| YNKMSL | MN833246.1 | MN870631.1 | MN861980.1 | 2019 | China | 36.040 | 104.220 | Tobacco |
| YNQJ | MN833245.1 | MN870639.1 | MN861977.1 | 2019 | China | 36.040 | 104.220 | Tobacco |
| YNrp | KM657120.1 | KM657117.1 | KM657114.1 | 2013 | China | 36.040 | 104.220 | Red Pepper |
| YNta | KM657121.1 | KM657118.1 | KM657115.1 | 2013 | China | 36.040 | 104.220 | Tobacco |
| Yongsang_Andong | OM022889.1 | OM022885.1 | OM022878.1 | 2021 | South Korea | 35.958 | 12.778 | Pepper |
| Yunnan | OM937130.1 | OM959369.1 | OM959368.1 | 2021 | China | 36.040 | 104.220 | *Glandularia* x *hybrida* |
| YY | MF159049.1 | MF159061.1 | MF159073.1 | 2016 | South Korea | 35.958 | 12.778 | Pepper |
| ZS | MT950318.1 | MT950317.1 | MT950316.1 | 2018 | China | 36.040 | 104.220 | *Perilla frutescens* |

**Table S2.** Putative major recombination sites found in genome segments of TSWV.

| Isolate name | Beginning Breakpoint | End breakpoint | Sampling location |
|---|---|---|---|
| **Segment L** | | | |
| MJ | 1 | 1133 | South Korea |
| HB | 75 | 1170 | South Korea |
| Tom-CA | 87 | 1464 | USA: California |
| TSWV-AS-P1 | 342 | 758 | South Korea |
| Tom-BL1 | 3382 | 3495 | USA: Texas |
| LS3 | 5166 | 5645 | South Korea |
| PepCal-22 | 8682 | 9029 | Italy |
| PepCal-24 | 8682 | 9029 | Italy |
| **Segment M** | | | |
| TSWV-LE | 1 | 2060 | China |
| 1-Ranunculus-2021 | 1002 | 2507 | South Korea |
| P1 | 1152 | 1570 | South Korea |
| Tom-CA | 1168 | 1576 | USA: California |
| Tarquinia | 1259 | 1513 | Italy |
| TSWV-BJFC-Ze | 4252 | 4979 | China |
| **Segment S** | | | |
| TswvTRPep | 131 | 3045 | Turkey |
| PepCal-22 | 1172 | 3242 | Italy |
| CG-1 | 1272 | 2393 | China |
| Tom-CA | 1621 | 2346 | USA: California |
| WA7 | 1718 | 1942 | Australia |
| Lazio17 | 2166 | 2491 | Italy |
| Pep-BL | 3176 | 3290 | USA: Texas |

**Table S3.** Percentage global similarity of proteins of TSWV compared with respective proteins of ANSV, GRSV, TCSV, CSNV and ZLCV.

| Protein | Group 1 | Group 2 | Similarity (%) |
|---|---|---|---|
| RdRp | TSWV | ANSV | 90 |
| | TSWV | CSNV | 89 |
| | TSWV | GRSV | 89 |
| | TSWV | TCSV | 89 |
| N | TSWV | ANSV | 79 |
| | TSWV | CSNV | 76 |
| | TSWV | GRSV | 80 |
| | TSWV | TCSV | 80 |
| | TSWV | ZLCV | 74 |
| NSs | TSWV | ANSV | 84 |
| | TSWV | CSNV | 86 |
| | TSWV | GRSV | 82 |
| | TSWV | TCSV | 81 |
| | TSWV | ZLCV | 73 |
| NSm | TSWV | ANSV | 88 |
| | TSWV | CSNV | 87 |
| | TSWV | GRSV | 85 |
| | TSWV | TCSV | 85 |
| | TSWV | ZLCV | 75 |
| Glycoprotein precursor | TSWV | ANSV | 87 |
| | TSWV | CSNV | 87 |
| | TSWV | GRSV | 81 |
| | TSWV | TCSV | 82 |
| | TSWV | ZLCV | 80 |

**Table S4**. Local similarity pairwise comparisons among TSWV and related viruses. Local similarity was calculated using a 90% threshold applied to the full-length protein sequence.

| Protein | Group 1 | Group 2 | Similarity ranges |
|---|---|---|---|
| RdRp | TSWV | ANSV | 100, 120, 140, 160, 180, 200, 220, 240, 260, 280, 300, 320, 340, 360, 380, 420, 500, 520, 580, 600, 620, 640, 660, 680, 1060, 1080, 1100, 1120, 1140, 1660, 1680, 1700, 1720, 1740, 1760, 1780, 1800, 1820, 1840, 1860, 1900, 1920, 1940, 1960, 2060, 2080, 2100, 2120, 2140, 2160, 2180, 2200, 2220, 2240, 2260, 2280, 2300, 2320, 2340, 2360, 2380, 2380, 2400, 2420, 2440, 2460, 2480, 2500, 2520, 2540, 2560, 2580, 2600, 2620, 2640, 2660, 2680, 2700, 2720, 2740, 2760 |
| | TSWV | CSNV | 400,440,460,480,540,560,980,1000,1020,1040,1320,1340,1360,1380, 1400,1420,1440,1460,1480,1600,1620,2820,2840 |
| | TSWV | GRSV | 700,720,740,760,780,800,820,840,860,880,900, 920,940,960,1160,1180, 1200,1220,1240,1260,1280,1300,1500,1520,1540,1560,1580,1640,1880, 2800 |
| | TSWV | TCSV | 1980,2000,2020,2040,2780 |
| N (nucleoprotein) | TSWV | GRSV | 50 |
| | TSWV | TCSV | 70 |
| | TSWV | ANSV | 90,110,130,150,170,190 |
| | TSWV | CSNV | 70,90 |
| NSs | TSWV | CSNV | 50,70,90,110,130,150,170,190,210,230,250,270, 290,310,330, 350,370,390,410 |
| | TSWV | TCSV | 210,230,250,270 |
| | TSWV | GRSV | 310 |
| | TSWV | ANSV | 50,70,90,110,130,190,230,250,270,290,310,330 |
| NSm | TSWV | ANSV | 50,70,90,230,250 |
| | TSWV | CSNV | 110,130,150,170,190,210 |
| | TSWV | ANSV | 90,110,130,150,170,190,210,230,250 |
| | TSWV | CSNV | 50,90,110,130,150,170,190,210,230,250 |
| Glycoprotein precursor | TSWV | CSNV | 100,120,140,160,180,200,220,240,500,520,540, 580,600,640 |
| | TSWV | ANSV | 260,280,300,320,340,360,380,400,420,440,460, 480,560,620, 660,680,700,720,740,760,780,800,820,840,860, 880,900,920, 940,960,980,1000,1020 |
| | TSWV | CSNV | 100,120,140,160,420,440,460,480,500,520,540, 560,580,600, 620,640,660,680,700,720,740,760 |
| | TSWV | ZLCV | 360,380,400,420,740 |

**Table S5**. Recombination events were identified in four proteins of TSWV and other tospoviruses. Group 1 and Group 2 represent different tospoviruses species that may have undergone recombination. Recombination positions were detected using the Proportion test implemented in SimPlot++ using grouped sequences (consensus).

| **Protein** | **Group 1** | **Group 2** | **start position** | **end position** | **score** |
|---|---|---|---|---|---|
| RdRp | TSWV | GRSV | 120 | 960 | 20.71 |
| | CSNV | TSWV | 1320 | 1500 | 12.39 |
| | ZLCV | TSWV | 1360 | 1540 | 12.28 |
| N | Not found | | | | |
| NSs | CSNV | TSWV | 270 | 310 | 4.93 |
| | TCSV | TSWV | 250 | 290 | 4.66 |
| | ANSV | TSWV | 270 | 330 | 4.54 |
| | TSWV | GRSV | 270 | 290 | 3.07 |
| NSm | CSNV | TSWV | 110 | 190 | 2.8 |
| | ANSV | TSWV | 110 | 190 | 2.77 |
| | TSWV | GRSV | 90 | 210 | 2.5 |
| | TSWV | TCSV | 90 | 210 | 2.49 |
| Glycoprotein precursor | TSWV | CSNV | 200 | 300 | 7.9 |
| | GRSV | TSWV | 620 | 700 | 6.24 |
| | TCSV | TSWV | 620 | 700 | 6.2 |
| | TSWV | ANSV | 200 | 420 | 4.15 |
