## Supplementary material for "Evolutionary origin and demographic trajectory of tomato spotted wilt virus (*Orthotospovirus tomatomaculae*)": Figure S1. Side-by-side comparison of phylogenetic trees reconstructed with whole genome sequences of TSWV segments L, M, and S. The blue-to-yellow co

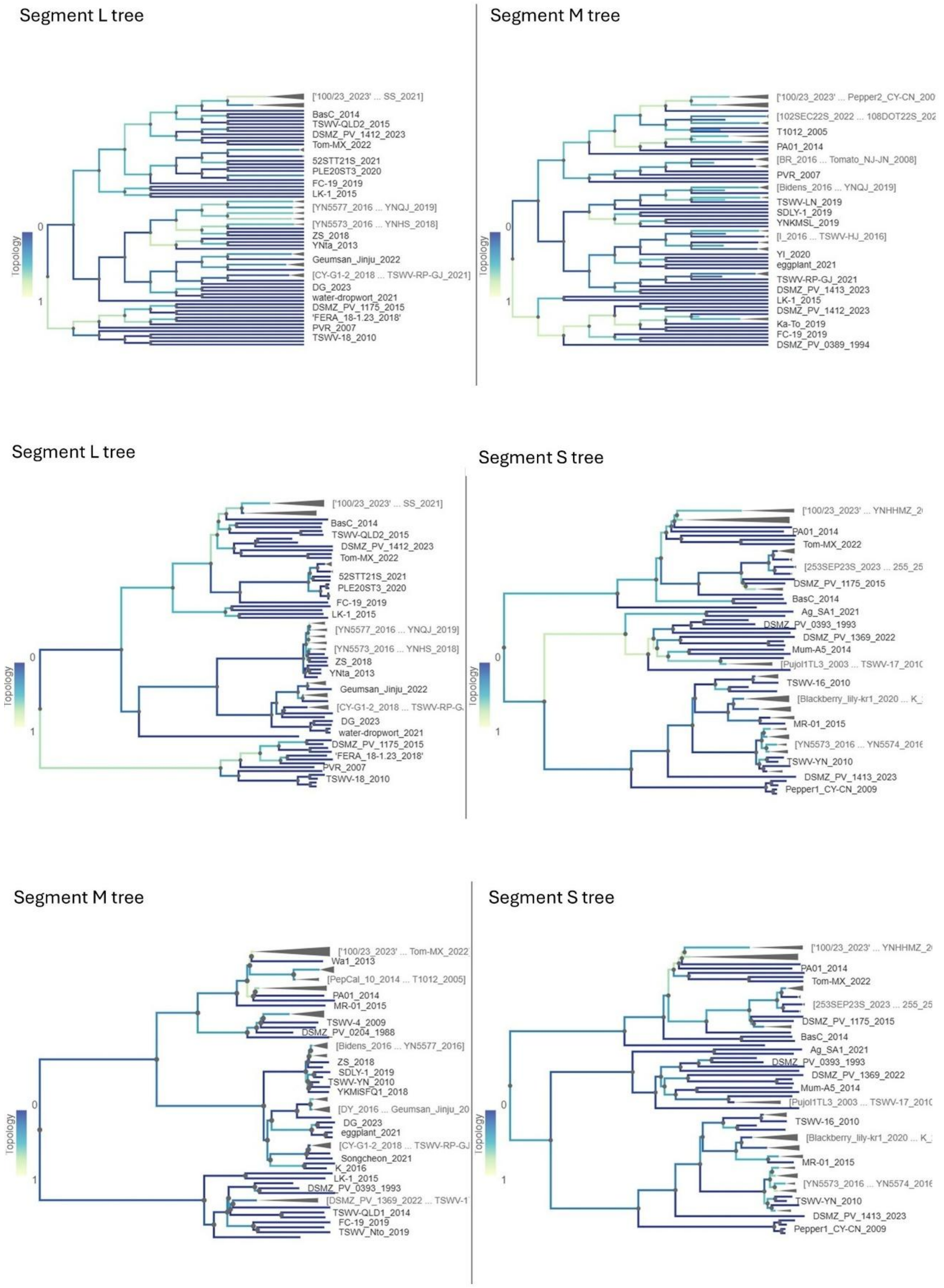


**Figure S1.** Side-by-side comparison of phylogenetic trees reconstructed with whole genome sequences of TSWV segments L, M, and S. The blue-to-yellow color gradient scheme indicates the similarity of best-matching branch patterns between the two trees. The Phylo.io tool (Robinson et al., 2016) was used for visualization.

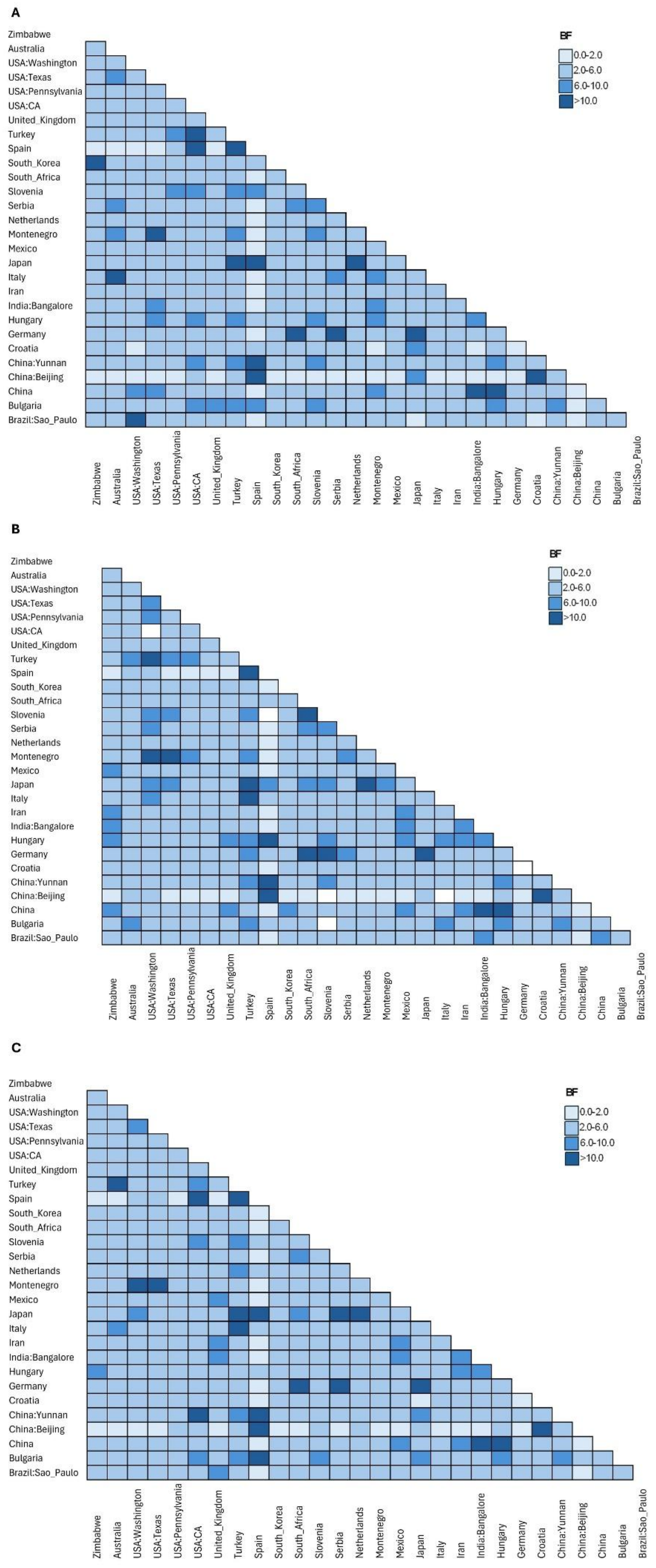
A
BF
0.0-2.0
2.0-6.0
6.0-10.0
>10.0
Zimbabwe
Australia
USA:Washington
USA:Texas
USA:Pennsylvania
USA:CA
United_Kingdom
Turkey
Spain
South_Korea
South_Africa
Slovenia
Serbia
Netherlands
Montenegro
Mexico
Japan
Italy
Iran
India:Bangalore
Hungary
Germany
Croatia
China:Yunnan
China:Beijing
China
Bulgaria
Brazil:Sao_Paulo
B
BF
0.0-2.0
2.0-6.0
6.0-10.0
>10.0
C
BF
0.0-2.0
2.0-6.0
6.0-10.0
>10.0

**Figure S2.** Bayes factor (BF) plots as the evidence of TSWV migration among countries according to phylogeographic analyses performed with sequence data of a) Segment L, B) Segment M and C) Segment S. The legend indicates BF values with the following interpretation: 0.0-2.0 no or weak evidence, 2.0-6.0 positive to strong evidence and >10 very strong evidence of TSWV migration between the indicated countries (Kass & Raftery, 1995).

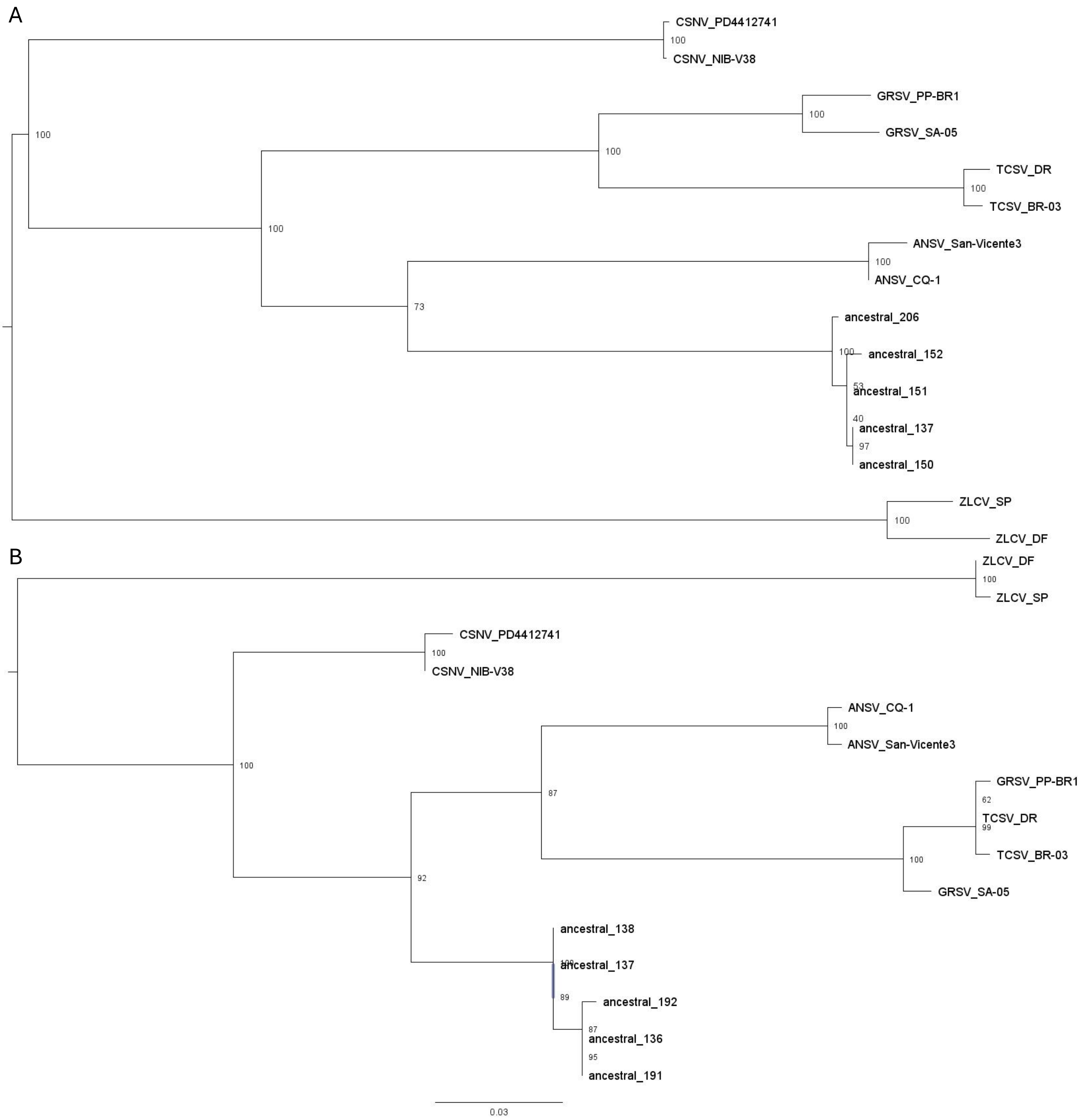

A
CSNV_PD4412741
100
CSNV_NIB-V38
GRSV_PP-BR1
100
GRSV_SA-05
100
TCSV_DR
100
TCSV_BR-03
100
100
ANSV_San-Vicente3
100
ANSV_CQ-1
73
ancestral_206
100
ancestral_152
53
ancestral_151
40
ancestral_137
97
ancestral_150
ZLCV_SP
100
ZLCV_DF
B
ZLCV_DF
100
ZLCV_SP
CSNV_PD4412741
100
CSNV_NIB-V38
ANSV_CQ-1
100
ANSV_San-Vicente3
100
GRSV_PP-BR1
62
TCSV_DR
99
87
100
TCSV_BR-03
GRSV_SA-05
92
ancestral_138
100
ancestral_137
89
ancestral_192
87
ancestral_136
95
ancestral_191
0.03

C

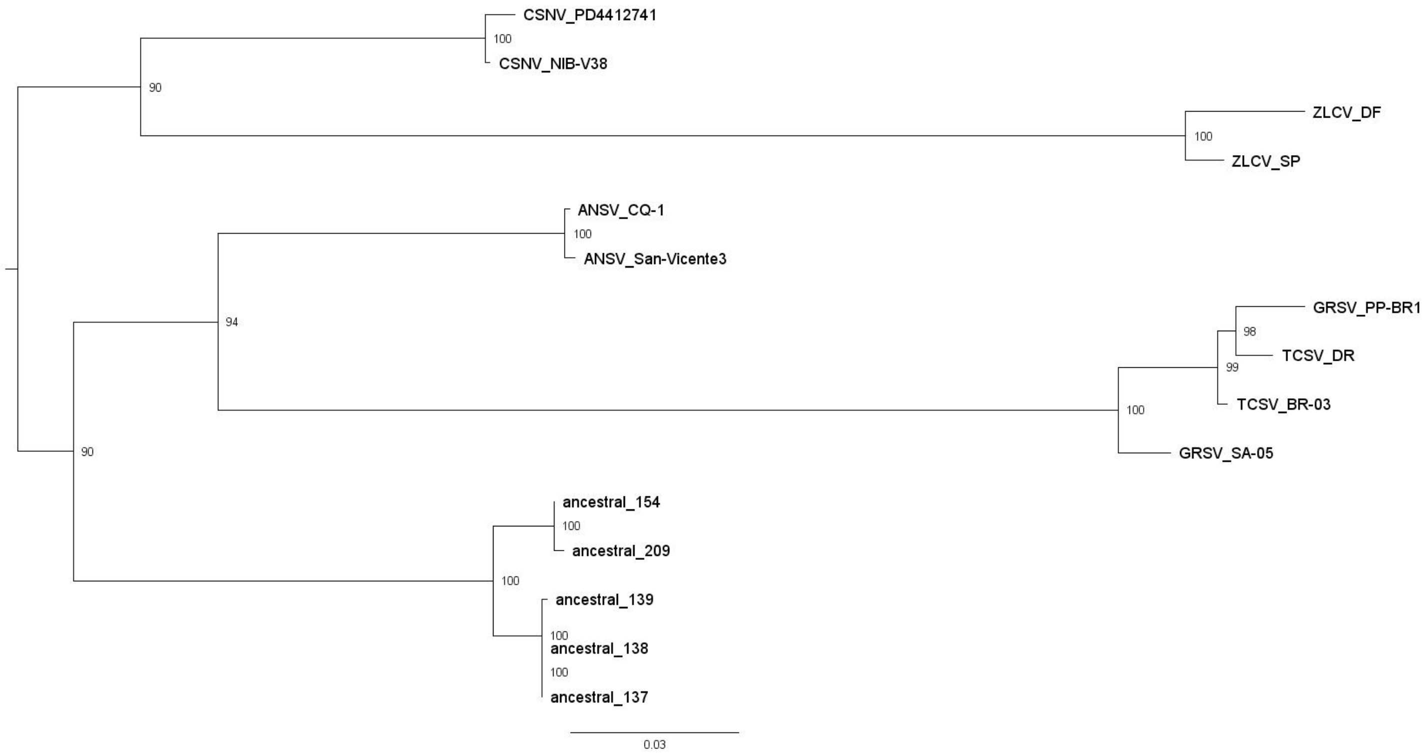
CSNV_PD4412741
100
CSNV_NIB-V38
90
ZLCV_DF
100
ZLCV_SP
ANSV_CQ-1
100
ANSV_San-Vicente3
94
GRSV_PP-BR1
98
TCSV_DR
99
TCSV_BR-03
100
GRSV_SA-05
90
ancestral_154
100
ancestral_209
100
ancestral_139
100
ancestral_138
100
ancestral_137
0.03

D

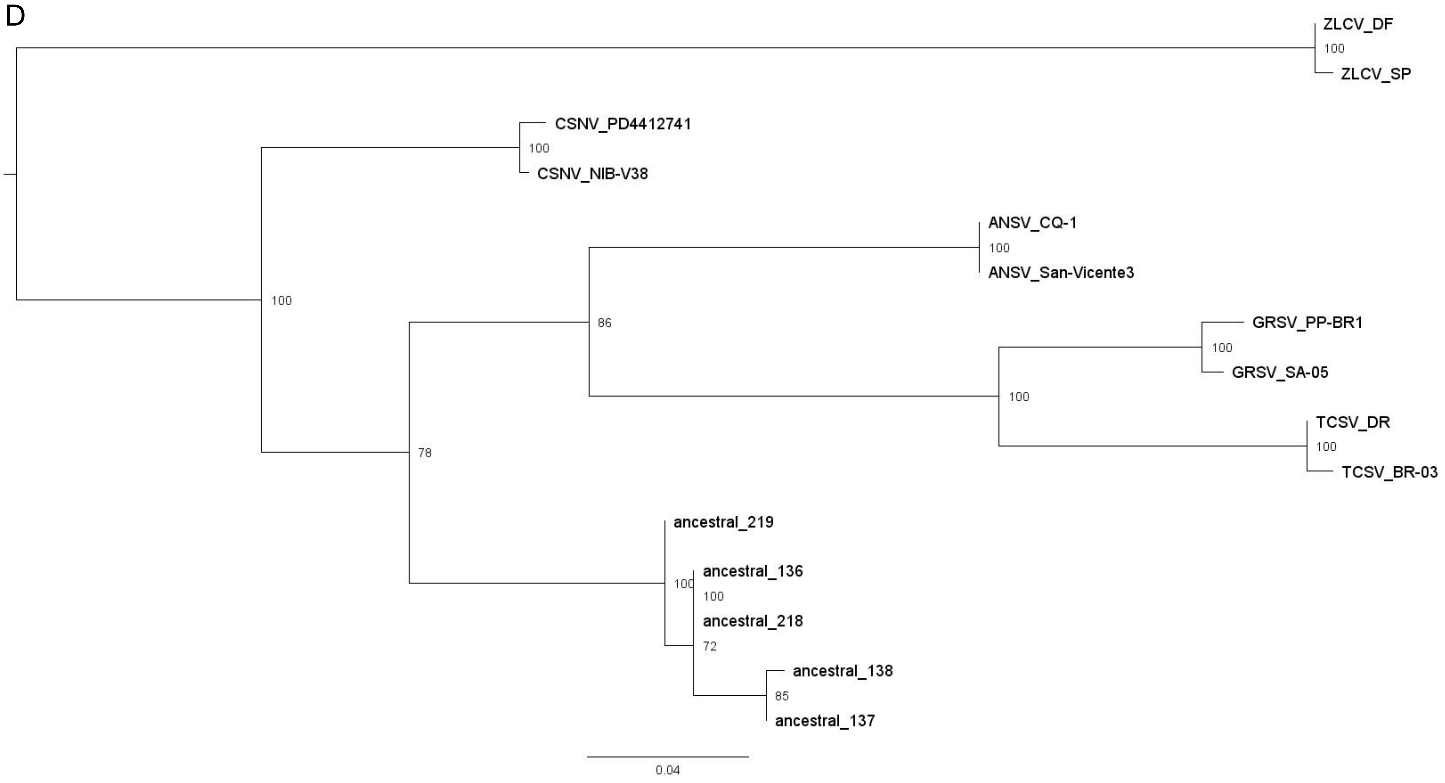
ZLCV_DF
100
ZLCV_SP
CSNV_PD4412741
100
CSNV_NIB-V38
100
ANSV_CQ-1
100
ANSV_San-Vicente3
86
GRSV_PP-BR1
100
GRSV_SA-05
100
TCSV_DR
100
TCSV_BR-03
78
ancestral_219
100
ancestral_136
100
ancestral_218
72
ancestral_138
85
ancestral_137
0.04

E

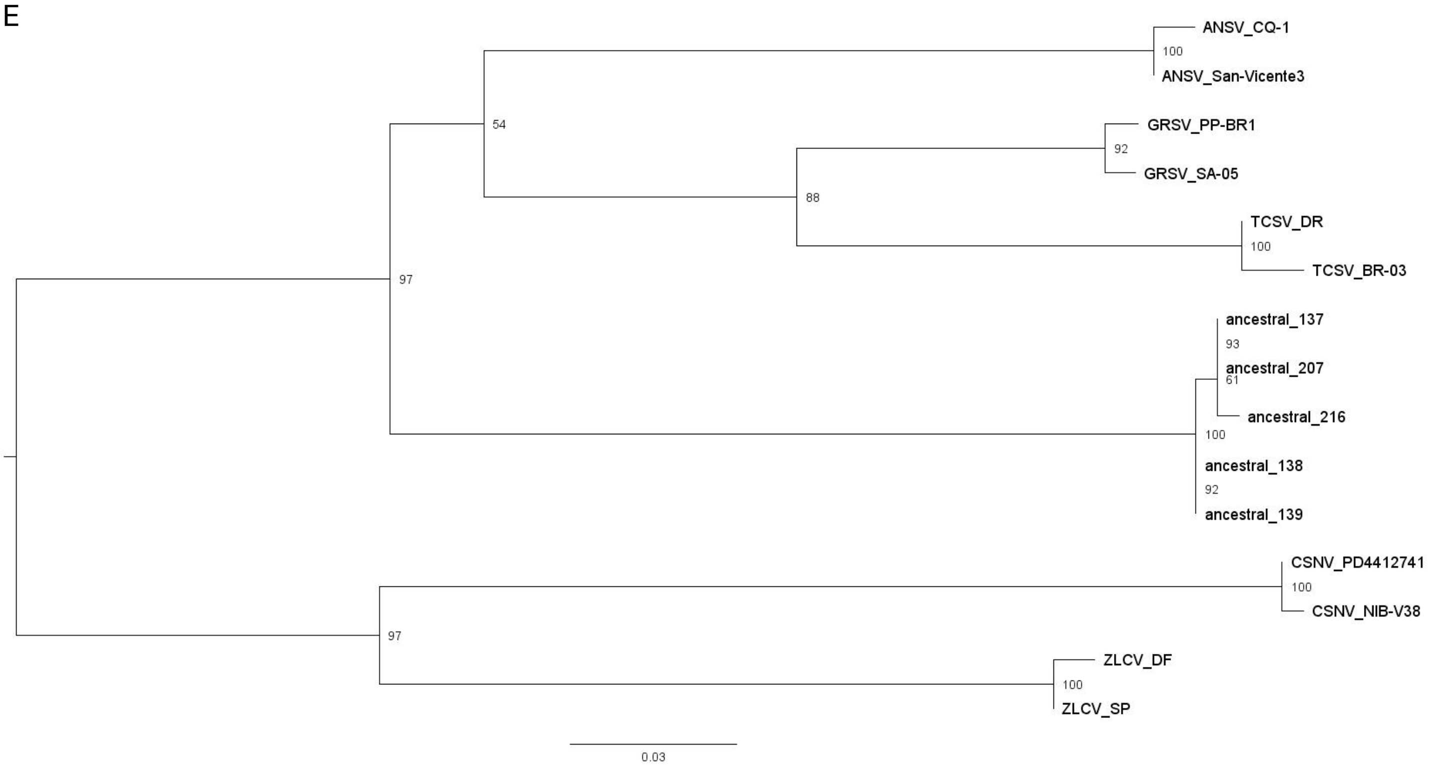


**Figure S3.** Maximum-likelihood trees of tospoviruses species reconstructed with A) RNA-dependent RNA polymerase (RdRp), B) movement protein (NSm), C) glycoprotein precursor, D) silencing suppressor (NSs), and E) nucleocapsid (N) protein sequences, using the IQTree program. Ancestral protein sequences were deduced from current sequences using the web server Fireprot ASR. Values on the nodes represent ultrafast bootstrap support, and the UFBoot trees were optimized by applying the nearest neighbor interchange directly to bootstrap alignments. The scale bar indicates branch length as the number of amino acid substitutions per site.
